# Exploring Antibiotic Resistance in Diverse Homologs of the Dihydrofolate Reductase Protein Family through Broad Mutational Scanning

**DOI:** 10.1101/2025.01.23.634126

**Authors:** Karl J. Romanowicz, Carmen Resnick, Samuel R. Hinton, Calin Plesa

## Abstract

Current antibiotic resistance studies often focus on individual protein variants, neglecting broader protein family dynamics. Dihydrofolate reductase (DHFR), a key antibiotic target, has been extensively studied using deep mutational scanning, yet resistance mechanisms across this diverse protein family remain poorly understood. Using DropSynth, a scalable gene synthesis platform, we designed a library of 1,536 synthetic DHFR homologs representing 778 species of bacteria, archaea, and viruses, including clinically relevant pathogens. A multiplexed *in vivo* assay tested their ability to restore metabolic function and confer trimethoprim resistance in an *E. coli* Δ*folA* strain. Over half of the synthetic homologs rescued the phenotype without supplementation, and mutants with up to five amino acid substitutions increased the rescue rate to 90%, highlighting DHFR’s evolutionary resilience. Broad Mutational Scanning (BMS) of homologs and 100,000 mutants provided critical insights into DHFR’s fitness landscape and resistance pathways, representing the most extensive analysis of homolog complementation and inhibitor tolerance to date and advancing our understanding of antibiotic resistance mechanisms.

**Graphical Abstract:** 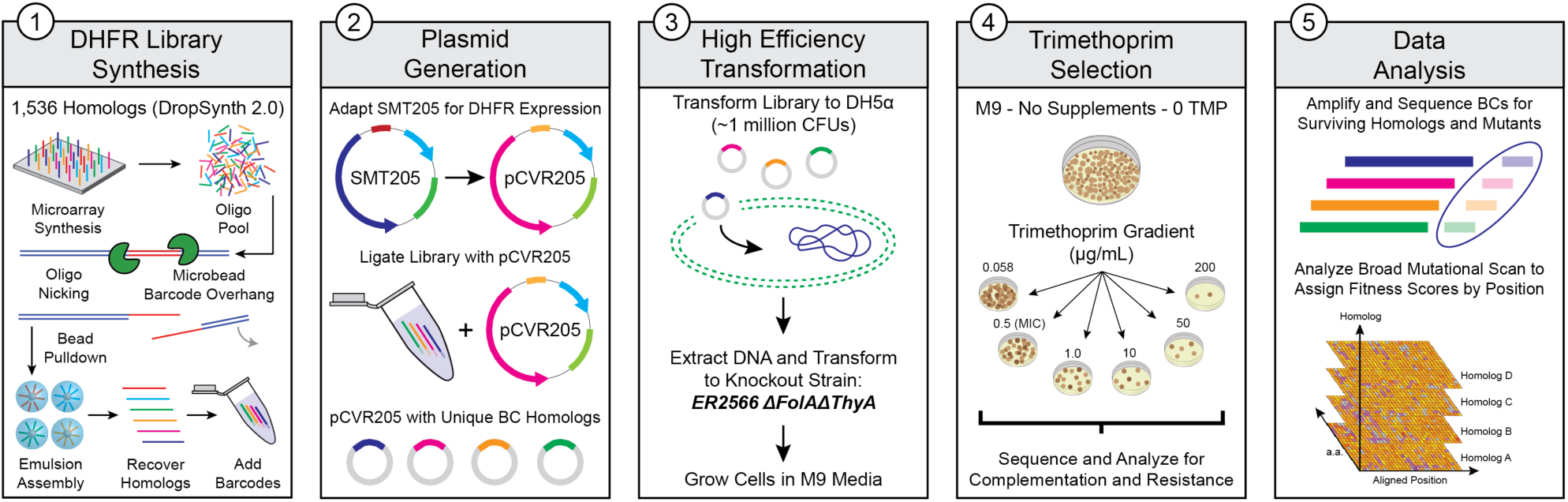

**Teaser:** DropSynth technology enables scalable and cost-effective exploration of antibiotic resistance across the DHFR protein family.

## Introduction

Antimicrobial resistance (AMR) is a pressing challenge in modern medicine, responsible for 4.95 million global deaths in 2019, surpassing the combined annual deaths from tuberculosis, malaria, and HIV/AIDS [1,2]. The extensive use of antibiotics in clinical and industrial settings has fueled the emergence of bacterial resistance mechanisms, jeopardizing the treatment of common infections [3]. Among antibiotic classes, dihydrofolate reductase (DHFR) inhibitors remain crucial for combating bacterial infections [4]. DHFR, a key enzyme in folate biosynthesis, reduces dihydrofolate to tetrahydrofolate, essential for DNA synthesis and cell proliferation [5]. Trimethoprim (TMP), a synthetic DHFR inhibitor, disrupts folate metabolism and inhibits bacterial growth [6]. However, increasing occurrences of bacterial resistance to TMP underscores the need for continued research into its effectiveness across diverse members of the DHFR protein family.

The DHFR protein family, essential for DNA synthesis across all organisms, has undergone significant divergent evolution, resulting in structural and functional variations with as little as 30% sequence identity between bacterial species [7]. This diversity complicates efforts to address antibiotic resistance, as it enables bacteria to develop various strategies to evade DHFR inhibition. These strategies include mutations in the DHFR gene that alter the enzyme’s structure and reduce drug binding affinity [8], acquisition of plasmid-encoded DHFR genes that produce enzymes with lower drug sensitivity [9,10], overexpression of DHFR to counteract the inhibitory effects of trimethoprim through increased enzyme abundance [11], and the development of alternative metabolic pathways that bypass DHFR activity [12,13]. The diversity of these resistance mechanisms highlights the adaptability of bacteria and the challenges in sustaining the efficacy of DHFR inhibitors such as trimethoprim as antibiotic treatments.

Given the critical role of DHFR in bacterial metabolism and the growing threat of antibiotic resistance, there is an urgent need to explore the sequence diversity within the DHFR family to understand trimethoprim efficacy and develop strategies to counteract resistance. Conventional methods like Deep Mutational Scanning (DMS) have been invaluable for revealing protein function and evolution by systematically examining how mutations affect functional landscapes, evolutionary constraints, and structure-function relationships [14,15,16]. However, DMS has been limited by existing gene synthesis techniques, restricting its analysis to single or few amino acid mutations in only a small number of proteins [17] and unable to probe the diversity of protein families that consist of thousands of members within a complex global landscape. This is due to several challenges faced by traditional gene synthesis techniques, including sequence complexity [18], length limitations imposed by phosphoramidite chemistry, which restricts oligonucleotide synthesis to fewer than 300 bases [19,20], and difficulties in assembling full-length genes from shorter fragments [21]. Additional hurdles include increasing error rates with longer sequences [22], limited high-throughput capacity [23], and high costs associated with large-scale gene libraries [24].

To address these challenges, we utilized DropSynth, a previously developed scalable and cost-effective method for constructing large, pooled libraries of systematically designed full-length gene sequences [25,26,27]. DropSynth enhances the traditional gene synthesis process through optimized oligonucleotide design, large-scale multiplexing, droplet-based emulsion assembly, large reductions in reagent volumes, and increased throughput, providing an efficient and cost-effective solution for the large-scale synthesis of gene-length sequences [25,26]. It works by compartmentalizing microarray-derived oligonucleotides into vortexed emulsions, where each water-in-oil droplet contains the oligonucleotides needed to assemble a single gene. This compartmentalization removes cross-hybridization between the fragments for different genes, allowing for proper full-length assembly for large numbers of genes in the same reaction. During this process, random mutations are introduced around the target sequences due to errors in oligonucleotide synthesis or DropSynth assembly. These mutations can be further analyzed to gain deeper insights into gene function [25]. Consequently, this approach generates vast libraries containing thousands of gene homologs and tens of thousands of mutant variants in parallel. Each variant is tagged with a random barcode, facilitating their transformation into bacterial knock-out models and subsequent recovery for functional assays, achieving a level of diversity that far exceeds conventional DMS experiments.

In this study, we combined DropSynth technology with Broad Mutational Scanning (BMS), an innovative approach for mapping the fitness landscape of diverse protein homologs generated through DropSynth-enabled gene synthesis [25]. Building on previous work where we designed and assembled two codon-optimized versions of a 1,536-member DHFR homolog library [26], we pursued four primary objectives: (**1**) to demonstrate the effective use of DropSynth-assembled DHFR libraries in complementing metabolic function in an *E. coli* knockout strain, (**2**) to evaluate the ability of DHFR homologs from diverse evolutionary lineages to confer resistance to the antibiotic trimethoprim, (**3**) to identify specific mutations in pathogenic DHFR variants that contribute to antibiotic resistance, and (**4**) to validate our multiplexed assay results using dial-out PCR [28]. To achieve these goals, we cloned the DHFR libraries into barcoded expression plasmids, transformed them into *E. coli ER2566* Δ*folA*Δ*thyA* cells [29], and screened for their ability to complement the *folA* knockout phenotype while testing resistance to trimethoprim across a concentration gradient. Dial-out PCR enabled independent testing of select homologs, allowing us to assess variant performance without competitive interference from other homologs to validate our pooled fitness results. This proof-of-concept study demonstrates the power of DropSynth-assembled libraries in multiplexed functional assays, showcasing our ability to construct and functionally characterize a diverse pool of DNA sequences for rational exploration of sequence-function relationships within the DHFR protein family at an unprecedented scale. It also provides a diverse dataset for training machine learning models in generative structure-function modeling. Ultimately, this work advances our understanding of the evolutionary mechanisms behind antibiotic resistance in the DHFR family, informing strategies to address this critical health challenge.

## Results

### DropSynth Assembly of DHFR Gene Homologs

We previously designed and assembled two codon versions of a 1,536-member library of dihydrofolate reductase (DHFR) homologs [26]. The Codon 1 library was generated using a weighted random codon assignment strategy, in which codon weights were based on their frequency in the *E. coli* genome, while excluding rare codons to enhance translational efficiency. In contrast, the Codon 2 library was constructed using a lookup table to systematically select codons that are maximally divergent from those in Codon 1, prioritizing sequence diversity at the expense of potential translational efficiency. These libraries were assembled using our top-performing polymerase, KAPA HiFi, barcoded with a random 20-nt assembly barcode, ligated into an expression plasmid (pCVR205), and transformed into DH5α cells to maximize ligation efficiency and library yield (**Fig. 1A**; see **Methods** for more details). Using our optimized protocol, we successfully assembled 1,208 unique homologs across both codon versions, representing a phylogenetically diverse collection of DHFR homologs with varying evolutionary distances from *E. coli* (**Fig. 1B**). These homologs encompass bacteria, archaea, and viruses, representing 19 phyla, 32 classes, 74 orders, 159 families, 330 genera, and 778 species, with most belonging to bacterial phyla such as Actinomycetota, Bacillota, Bacteroidota, Mycoplasmatota, and classes of Pseudomonadota (**Table S1**). Our library design intentionally incorporates numerous pathogenic DHFR homologs (**Fig. 1B**), including ESKAPE bacterial pathogens [30,31].

**Figure 1.**
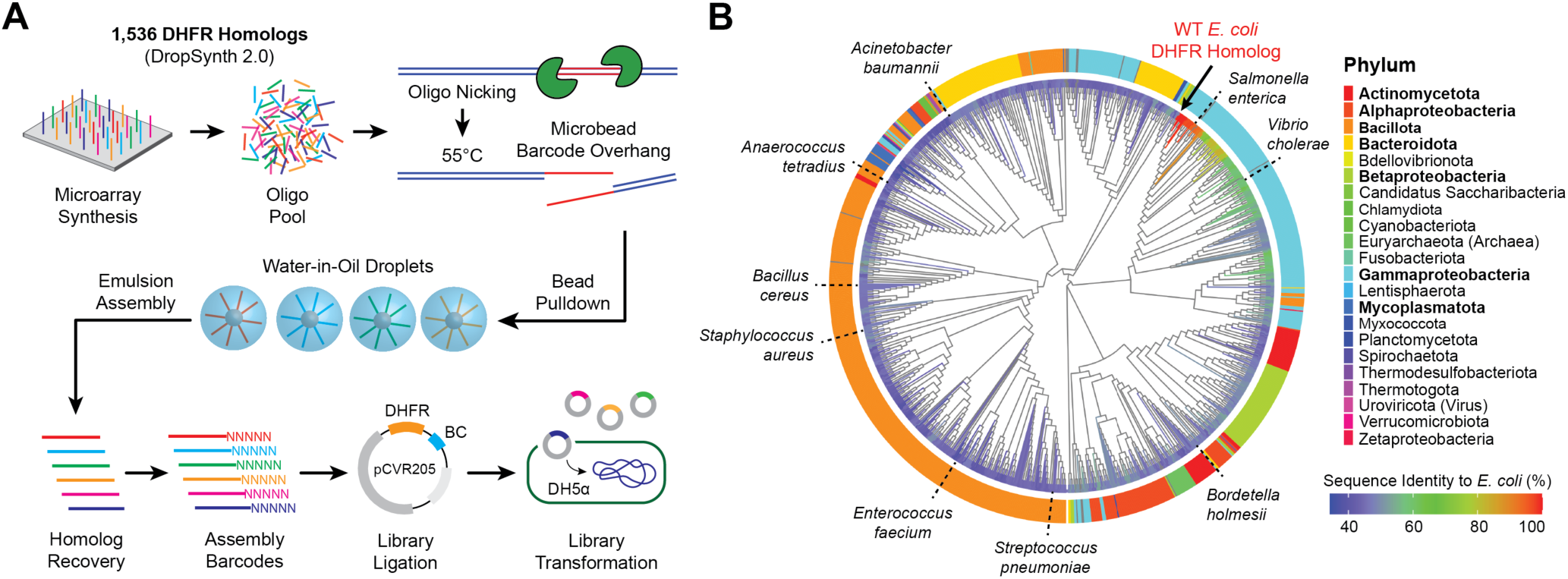
DropSynth Assembly of 1,536-DHFR Gene Homologs. (**A**) Schematic of the DropSynth 2.0 process: microarray-derived oligos are amplified and pulled down by barcoded beads that selectively capture oligos for gene assembly. Beads are emulsified in droplets with genes assembled via polymerase cycling assembly (PCA). After breaking the emulsion, genes are barcoded, ligated into an expression plasmid, and transformed into DH5α cells. (**B**) Maximum likelihood phylogenetic tree of 1,208 successfully assembled DHFR homologs from the 1,536-member library. Branch color indicates percent sequence identity to WT *E. coli* DHFR, with taxonomic diversity annotated at the class level for the phylum Pseudomonadota. Known pathogenic homologs are labeled around the tree for reference. Phyla in **bold** represent the highest unique sequence abundances in the DHFR library.

### Complementation Assay of DHFR Homologs and Mutants

Our first objective was to demonstrate how DropSynth-assembled libraries can be seamlessly integrated into multiplex functional assays. Here, we tested our assembled DHFR homologs, derived from diverse evolutionary backgrounds, for their ability to rescue a *folA* knockout phenotype in *E. coli*. The barcoded DHFR homologs under a T7 promoter in the pCVR205 expression plasmid were transformed into folate- and thymidine-deficient *E. coli ER2566* Δ*folA*Δ*thyA* cells and grown on LB agar with chloramphenicol and thymidine, followed by growth on supplemented M9 media. The surviving cells were screened for complementation by growing them on non-supplemented M9 media without thymidine at 37°C for 22 hours (**Fig. 2A**). This auxotrophic strain, derived from the ER2566 background [32], requires external thymidine for growth and is commonly used to express recombinant DHFR proteins without interference from the host’s native DHFR activity. Notably, the *thyA* gene was knocked out of the genome and placed on the expression plasmid to prevent metabolic toxicity effects associated with DHFR overexpression [33]. The complementation assay on non-supplemented M9 media tests the cell’s ability to survive under stringent conditions where reductase activity depends solely on a single DHFR gene from our DropSynth-assembled library. Therefore, changes in the fraction of barcodes recovered after growth without thymidine supplementation reflect the extent of DHFR functional complementation. This assay is essential for establishing a baseline to compare barcode counts across supplemented, non-supplemented, and trimethoprim-inhibited conditions (see **Methods** for more details).

**Figure 2.**
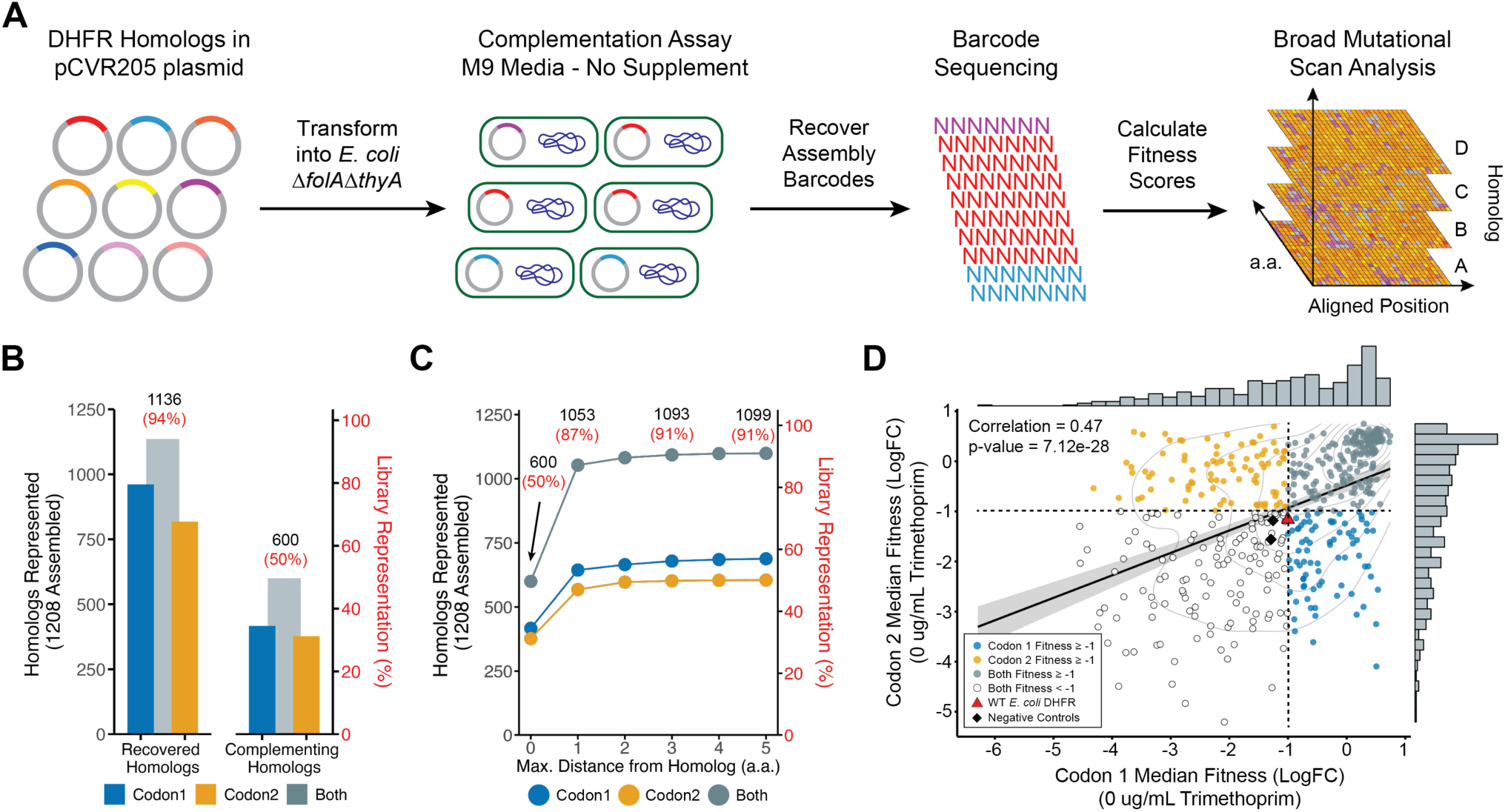
Complementation Assay of DHFR Homologs and Mutants. (**A**) Barcoded DHFR homologs in the pCVR205 plasmid were transformed into folate-deficient *E. coli ER2566* Δ*folA*Δ*thyA* cells using a pooled complementation assay with growth on solid media. Fitness scores were based on the Log_2_ fold-change in barcode recovery between supplemented and non-supplemented conditions. Sequence alignment enabled mapping fitness onto the wild-type (WT) *E. coli* DHFR sequence. (**B**) Of the 1,208 DropSynth-assembled homologs, we recovered 961 from Codon 1 and 818 from Codon 2 with a minimum of 10 sequences per barcode, with 1,136 homologs represented across both codon libraries under complementation conditions (94% total library coverage). Selecting homologs showing some degree of complementation (fitness ≥ −1) revealed 416 homologs from Codon 1 (34%) and 377 homologs from Codon 2 (31%), totaling 600 unique homologs capable of complementation across both codon versions, achieving 50% coverage of the total library. (**C**) Adding mutant assemblies (within 5 amino acid distance), we recovered 1,099 homologs capable of complementation (fitness ≥ −1), achieving 91% of total library coverage. (**D**) Fitness values for 485 homologs shared between codon versions showed a significant correlation (ρ = 0.47; Pearson), with points representing fitness in Codon 1 (blue), Codon 2 (orange), or both (teal). Uncolored points represent homologs with low fitness during complementation (fitness < −1). WT *E. coli* DHFR and negative controls (D27N, mCherry) are plotted for reference. Dashed lines indicate the minimum fitness threshold for complementation.

Barcodes were previously mapped to their corresponding DHFR variants using Illumina MiSeq sequencing. Illumina NovaSeq sequencing of the assembly barcodes recovered from all nine experimental growth conditions (LB media, supplemented M9 media, non-supplemented M9 media, and six trimethoprim concentrations in non-supplemented M9 media) generated 1,010,919,383 sequence reads from both codon version libraries. These reads corresponded to 1,136,880 unique assembly barcodes, each with at least one read count (**Table S2**). Under the non-supplemented condition (i.e., complementation), we recovered 122,525,216 reads corresponding to 403,154 unique barcodes, of which 227,451 (56.4%) were present in the mapping data, after applying a minimum threshold of 10 sequence reads per barcode (**Fig. S1**). Among these mapped barcodes, 65% (147,868) corresponded to perfect assemblies, while 35% (79,617) were associated with mutant variants. In total, the mapped barcodes corresponded to 1,136 unique homologs across both codon versions, representing 94% of the 1,208 successfully assembled homologs recovered from the original library of 1,536 designs (**Fig. 2B; Table S3**). A notable difference between the codon versions was the substantially higher proportion of mutant assemblies with over 100 amino acid substitutions in Codon 2 (10.3%) compared to Codon 1 (2.2%), which contributed to a lower recovery of perfect assemblies in Codon 2 (75.7%) versus Codon 1 (84.3%) (see **Fig. S1** for details).

After filtering for perfect assembly homologs with at least 5 unique barcodes, we retained 797 homologs from Codon 1 and 666 homologs from Codon 2 under the complementation conditions, representing 970 unique homologs across both codon versions (**Table S3**). We also retained 59,763 unique mutant variants associated with the perfect assembly homologs from Codon 1 and 49,691 from Codon 2, representing 109,454 unique mutant variants across both codon versions. Of these, 12,274 unique mutants from Codon 1 and 16,060 from Codon 2, were within 5 amino acids of the perfect assembly homologs, representing 28,334 high-quality mutants. Each DHFR homolog had a median of 18 unique mutant variants for both codon versions. We calculated fitness scores associated with each DHFR homolog and its mutant variants based on the log_2_ fold-change in barcode recovery between M9 supplemented and non-supplemented media conditions (**Fig. 2A**).

Among the perfect assembly homologs (with at least 5 unique barcodes) that showed some degree of complementation (i.e., fitness ≥ −1), we identified 416 homologs from Codon 1 and 377 homologs from Codon 2. Together, these accounted for 600 unique homologs across both codon versions, representing 50% of the 1,208 DropSynth-assembled library (**Fig. 2B**). By including high-quality mutant assemblies with up to five amino acid differences from their designed homologs, we recovered 1,099 unique homolog-centered-clusters capable of complementation (fitness ≥ −1) across both codon versions, achieving 91% coverage of the library members passing quality checks under complementation conditions (**Fig. 2C**). Most of these gain-of-function mutants resulted from single amino acid substitutions, totaling 1,053 unique homologs across both codon versions. False-positive rates (fitness ≥ −1), estimated using highly mutated variants (>50 amino acid substitutions) with a minimum of 5 unique barcodes, were 0.6% in the Codon 1 library and 1.3% in the Codon 2 library at complementation (**Table S3**). Mutant assemblies with more than 5 amino acid changes from the designed homologs were excluded from downstream analyses to ensure fitness effects accurately reflected the intended sequences rather than artifacts from excessive mutations.

Of the 970 perfect assembly homologs retained across both codon versions under the complementation conditions, 493 were shared between codon libraries (**Table S4**). The fitness values of the shared homologs exhibited a significant positive correlation (ρ = 0.47, Pearson; *P* = 7.12 × 10⁻²⁸; **Fig. 2D**). Notably, most of the complementing homologs (fitness ≥ −1) were common to both codon libraries, totaling 202 homologs (41% shared; **Fig. 2D**). An additional portion of complementing homologs were codon-specific, with 77 homologs complementing only with the Codon 1 optimized sequence design (16% shared) and 86 homologs complementing only with the Codon 2 design (17% shared) (**Fig. 2D**). The remaining 128 shared homologs (26%) were incapable of complementation (fitness < −1) (**Fig. 2D**). The wild-type (WT) *E. coli* DHFR homolog, spiked into both libraries as a positive control, showed consistent fitness across codon versions (**Fig. 2D**). Likewise, the low-function D27N DHFR variant [34] and non-catalytic mCherry protein [35], spiked as negative controls (see **Methods** for more details), exhibited consistent fitness (**Fig. 2D**).

The fitness distribution of perfect assembly homologs under complementation conditions falls into two categories: homologs capable of complementation (fitness ≥ −1), which were used for Broad Mutational Scanning (BMS), and dropout homologs (fitness < −1), which were used for gain-of-function (GOF) analysis (**Fig. 3A**). We focused on homologs from the Codon 1 library, optimized for expression in *E. coli*, for both analyses. Among the 797 homologs retained from Codon 1 under the complementation conditions, we identified 416 capable of complementation and 381 dropouts (**Fig. 3A**). Complementing and dropout homologs are evenly distributed across the phylogenetic tree, with some minor clustering within certain clades (**Fig. 3B**). Despite a median sequence identity of 46% (see **Fig. 1B** for reference), most of the distant homologs still complement the function of the native *E. coli* DHFR (**Fig. 3B; Fig. S2**). Several factors may contribute to the low fitness observed in dropout homologs, such as improper protein folding, metabolic mismatches with *thyA*, low *folA* gene expression leading to insufficient metabolic flux, undesired protein-protein interactions, or gene dosage toxicity due to excessively high *folA* expression, each of which can impact survival in the *E. coli* knockout model [33,36].

**Figure 3.**
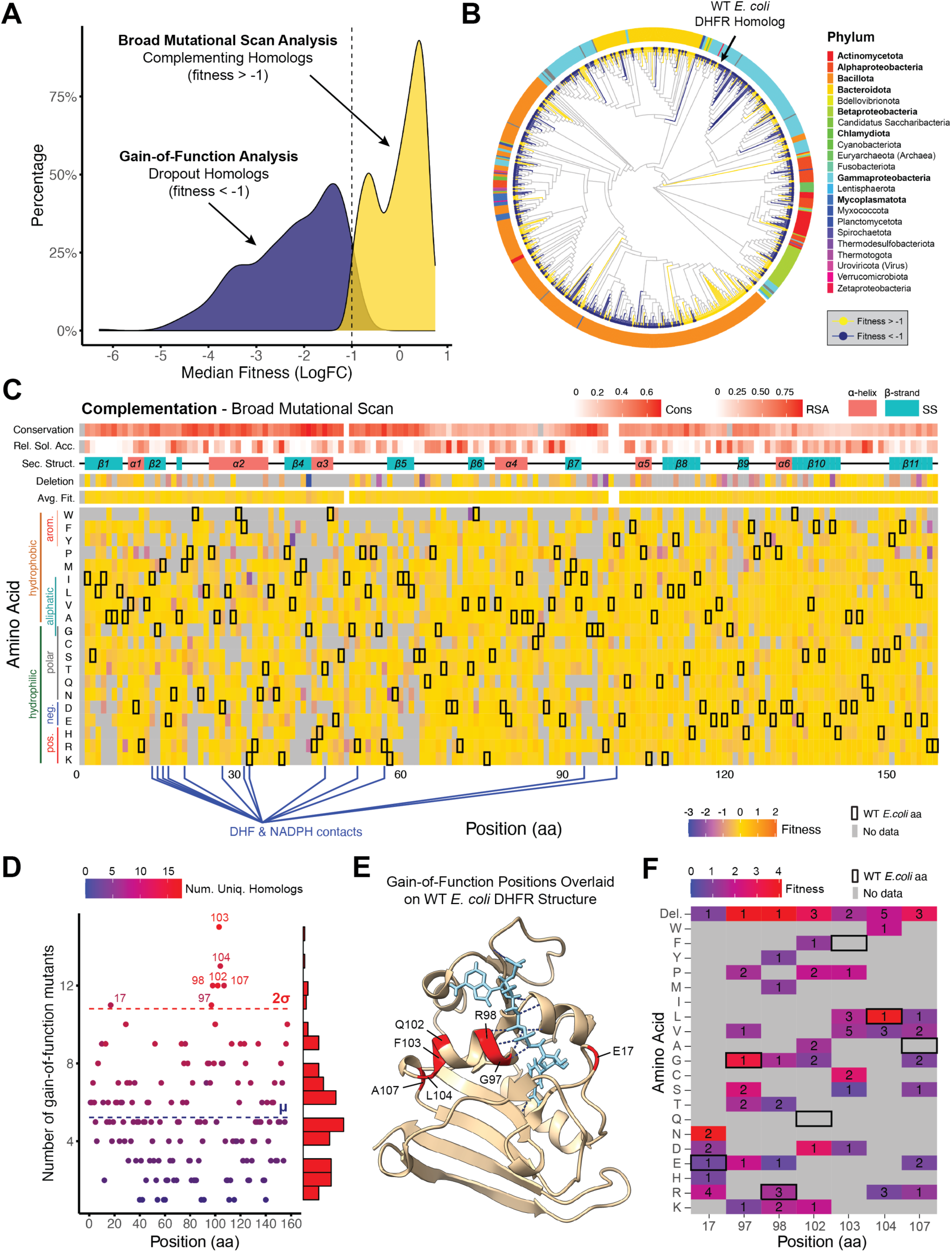
Broad Mutational Scanning and Gain-of-Function Mutations in Complementation Assay. (**A**) Fitness distribution of 797 DHFR homologs from the Codon 1 complementation assay, with 416 complementing homologs (fitness ≥ −1, yellow) used in BMS and 381 dropout homologs (fitness < −1, blue) used in GOF analysis. (**B**) Phylogenetic tree of 797 DHFR homologs, highlighting complementing and dropout homologs, with Pseudomonadota annotated at the class level. (**C**) Fitness landscape of 416 complementing homologs and 5,419 associated mutants within 5 amino acids mapped onto the WT *E. coli* DHFR sequence, showing average fitness by position and additional metrics (secondary structure, solvent accessibility, site conservation). (**D**) GOF analysis identified 476 mutants (fitness ≥ −1) associated with 196 dropout homologs, projected onto the WT *E. coli* DHFR sequence, highlighting seven statistically significant positions (residues 17, 97, 98, 102, 103, 104, and 107). (**E**) WT *E. coli* DHFR structure (4KJK) with GOF residues shaded in red, NADPH cofactor in cyan, and H^+^ bonds in blue. (**F**) Mean fitness and mutant frequency at each GOF position.

For the Broad Mutational Scanning (BMS) analysis, we selected all 416 homologs from the Codon 1 library that showed some degree of complementation (fitness ≥ −1) along with their 5,419 mapped mutants within a five-amino-acid distance. We then performed a multiple sequence alignment to identify equivalent residue positions. For each amino acid and position, we calculated the median fitness score across all homologs and mutants. The three-dimensional homolog data (**Fig. 2A right end**) was collapsed and projected onto the *E. coli* DHFR sequence (**Fig. 3C**), generating insights akin to those from conventional DMS approaches [37,38]. Average BMS fitness was significantly constrained at highly conserved sites (ρ = −0.51, Pearson; *P* = 6.54 × 10⁻⁹), indicating strong mutational constraints (**Fig. S3A**). In contrast, buried residues showed only a slight reduction in average fitness compared to solvent-accessible residues (ρ = 0.24, Pearson; *P* = 0.003), based on DSSP analysis of the 1H1T crystal structure (**Fig. S3B**). Higher mutational coverage is associated with increased positional fitness estimates (ρ = 0.49, Pearson; *P* = 8.63 × 10⁻¹¹). This correlation occurs because low-fitness homologs and mutants were more often excluded, leading to a bias that elevated the average positional fitness values (**Fig. S3C**). Notably, average BMS fitness did not differ significantly across secondary structures (helices, loops, strands; **Fig. S3D**).

In the gain-of-function (GOF) analysis, we identified 476 GOF mutants (fitness ≥ −1) with single amino acid substitutions associated with 196 of the 381 dropout homologs (fitness < −1). By aligning these mutations to the WT *E. coli* DHFR sequence, we identified seven statistically significant residues (positions 17, 97, 98, 102, 103, 104, and 107) based on GOF mutation frequency at each site (**Fig. 3D**). When mapped onto the *E. coli* DHFR protein structure (4KJK, Protein Data Bank) [39], all but one of these residues form a cluster in a small alpha-helix region (**Fig. 3E**). Within this region, residues at positions 97 and 98 appear to contribute to the binding pocket for nicotinamide adenine dinucleotide phosphate (NADPH), the essential cofactor for dihydrofolate reduction. Specifically, these residues form hydrogen bonds with the nicotinamide ring, helping to position the cofactor for catalysis (**Fig. 3E**). Additionally, the significant residue at position 17, within the critical Met20 loop (residues 17–20), plays a crucial role in stabilizing substrate and cofactor binding, undergoing conformational shifts during catalysis to align ligands in the active site for efficient dihydrofolate reduction [40]. At each significant GOF position, multiple unique single amino acid substitutions restored dropout homolog fitness, with deletions at certain positions yielding the highest fitness gains (**Fig. 3F**).

### Trimethoprim Resistance of DHFR Homologs and Mutants

Following complementation, our next objective was to test whether DropSynth-assembled libraries could maintain metabolic function under trimethoprim (TMP) exposure. To assess fitness, *E. coli ER2566* Δ*folA*Δ*thyA* cells containing barcoded DHFR plasmids were exposed to a trimethoprim gradient ranging from 0 to 400× the minimal inhibitory concentration (MIC) of 0.5 µg/mL in minimal M9 media. TMP concentrations included 0 (complementation), 0.058, 0.5 (MIC), 1.0, 10, 50, and 200 (400× MIC) µg/mL (**Fig. 4A**). Plasmid barcodes from cells surviving each TMP condition were recovered, amplified, and sequenced to quantify the abundance of each unique DHFR variant across the TMP gradient. This approach enabled us to identify high-fitness variants that thrived under increasing antibiotic pressure and low-fitness variants that were progressively depleted with rising trimethoprim concentrations.

**Figure 4.**
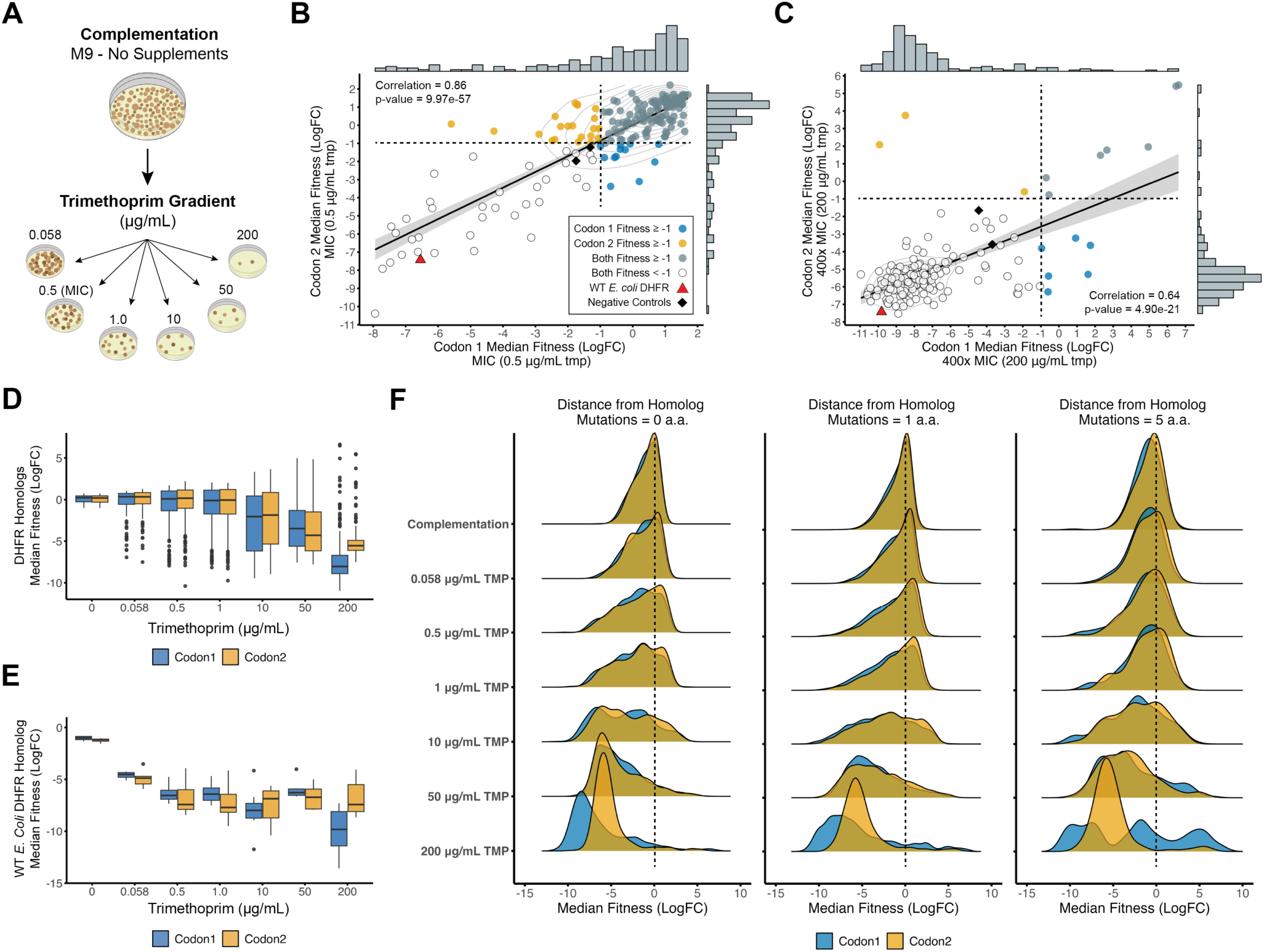
Trimethoprim Fitness Assay of DHFR Homologs and Mutants. (**A**) Overview showing how *E. coli ER2566* Δ*folA*Δ*thyA* cells containing unique barcoded DHFR homologs were grown across a trimethoprim gradient. Fitness scores were calculated based on the Log_2_ fold-change in barcode sequence recovery between trimethoprim-treated and supplemented conditions. (**B**) At MIC (0.5 µg/mL), fitness values for 190 shared homologs showed a strong positive correlation (ρ = 0.86; Pearson) between codon versions, with many homologs demonstrating resistance (fitness ≥ −1). (**C**) At 400× MIC (200 µg/mL), fitness values for 174 shared homologs showed a significant positive correlation (ρ = 0.64; Pearson) between codon versions, though most homologs could not maintain function (fitness < −1). Points are colored based on fitness for codon-specific homologs, shared homologs, or those with low fitness. The WT *E. coli* DHFR homolog and negative controls are plotted for reference. Dashed lines denote the minimum fitness thresholds for trimethoprim resistance. (**D**) Boxplot showing homolog fitness declined consistently with increasing TMP concentration across the gradient for both codon versions. (**E**) Boxplot showing WT *E. coli* DHFR homolog (positive control) fitness declined similarly across both codon versions. (**F**) Ridge plots show that the TMP gradient impacted the median fitness distributions for perfect assemblies and mutants with 1 or 5 amino acid differences, with survival decreasing as TMP concentration increased.

Among the 600 perfect assembly homologs that complemented across both codon versions (fitness ≥ −1; see **Fig. 2B** for reference), only 190 shared homologs were recovered at the MIC level. These shared homologs showed a significant positive fitness correlation (ρ = 0.86, Pearson; *P* = 9.97 × 10^-57^), with most maintaining metabolic function (fitness ≥ −1) at MIC selection (**Fig. 4B**). At 400× MIC (200 μg/mL), we recovered 174 shared homologs that displayed a positive but lower fitness correlation than at MIC (ρ = 0.64, Pearson; *P* = 4.90 × 10^-21^), with most unable to maintain function (fitness < −1) (**Fig. 4C**). Notably, we identified 7 shared homologs and 9 additional codon-specific homologs capable of maintaining function (fitness ≥ −1) under high TMP, indicating resistance to trimethoprim (**Fig. 4C**). These include members of Bacillota (e.g., *Clostridium* spp.), Bacteroidota (*Galbibacter marinus, Niabella soli*), and Pseudomonadota (e.g., *Neisseria flavescens*) (**Table S5**). We also recovered homologs with higher median fitness (max. fitness = 6.6) than observed in the complementation condition (max. fitness = 0.7; see **Fig. 2D**). This is likely due to the compositional nature of the data: as many variants die off, the relative fraction of tolerant variants increases, resulting in higher enrichment scores. Pearson fitness correlations for homologs shared between codon versions under each TMP condition are provided in Supplemental Materials (see **Fig. S4**).

Across the TMP gradient, increasing antibiotic concentrations consistently decreased median homolog fitness for both codon version libraries (**Fig. 4D**). This trend demonstrates that even high-fitness homologs experience reduced viability with increasing trimethoprim inhibition. The WT *E. coli* DHFR homolog, spiked into each library as a positive control, exhibited a similar pattern, with a steady fitness decline across the TMP gradient, highlighting the substantial metabolic stress imposed by trimethoprim (**Fig. 4E**). The fitness responses of the negative controls, D27N and mCherry, are presented alongside the WT *E. coli* DHFR homolog for comparison (see **Fig. S5**). When examining DHFR variants with either a single amino acid change or up to five amino acid changes, we observed declining fitness values with increasing TMP concentration that were comparable to those of the perfect assembly variants (i.e., 0 mutations), consistent across both codon versions (**Fig. 4F**). However, the ridge plots reveal a nuanced, codon-specific response at the highest TMP concentration (200 μg/mL). Homologs with up to five amino acid differences in the Codon 1 library showed significantly divergent fitness outcomes: where some displayed extremely low fitness (< −5), others experienced moderate declines (fitness between −1 and −5), and a subset achieved strong positive fitness (> 5). This variation suggests that specific protein mutations in the Codon 1 library may provide certain homologs with enhanced resistance to high TMP inhibition, indicating potential adaptive advantages unique to this codon version (see **Fig. S6** for more details).

To analyze the mutational fitness landscape across the TMP gradient, we again focused on homologs from the Codon 1 library optimized for *E. coli* expression. Of the 416 homologs capable of complementation under non-supplemented conditions (fitness ≥ −1; see **Fig. 3A** for reference), we observed a decline in the number of homologs resistant to trimethoprim inhibition as the antibiotic concentration increased across the gradient (see **Fig. S7**). Specifically, 318 homologs (76%) were resistant (fitness ≥ −1) at 0.058 µg/mL TMP, 246 homologs (59%) at 0.5 µg/mL (MIC), 226 homologs (54%) at 1.0 µg/mL, 128 homologs (31%) at 10 µg/mL, 80 homologs (19%) at 50 µg/mL, and only 27 homologs (7%) remained resistant at 200 µg/mL TMP (**Table S6**). Although most homologs were resistant to trimethoprim at the MIC level (59%), the absolute differences in fitness values between homolog pairs were not evenly distributed across the phylogenetic tree, exhibiting significant variation based on evolutionary distance (**Fig. 5A**). A Wilcoxon rank-sum test indicated a highly significant difference in fitness variation (W = 3.2 × 10^8^, *P* < 2.2 × 10⁻¹⁶), supported by Welch’s t-test, which showed a lower mean absolute fitness difference for closely related homologs compared to distantly related homologs (t = −63.5, df = 7,020, *P* < 2.2 × 10⁻¹⁶; **Table S7**). These results suggest that evolutionary relatedness significantly influences homolog fitness similarity under MIC selection, with closely related species tending to exhibit more similar fitness responses to trimethoprim treatment.

**Figure 5.**
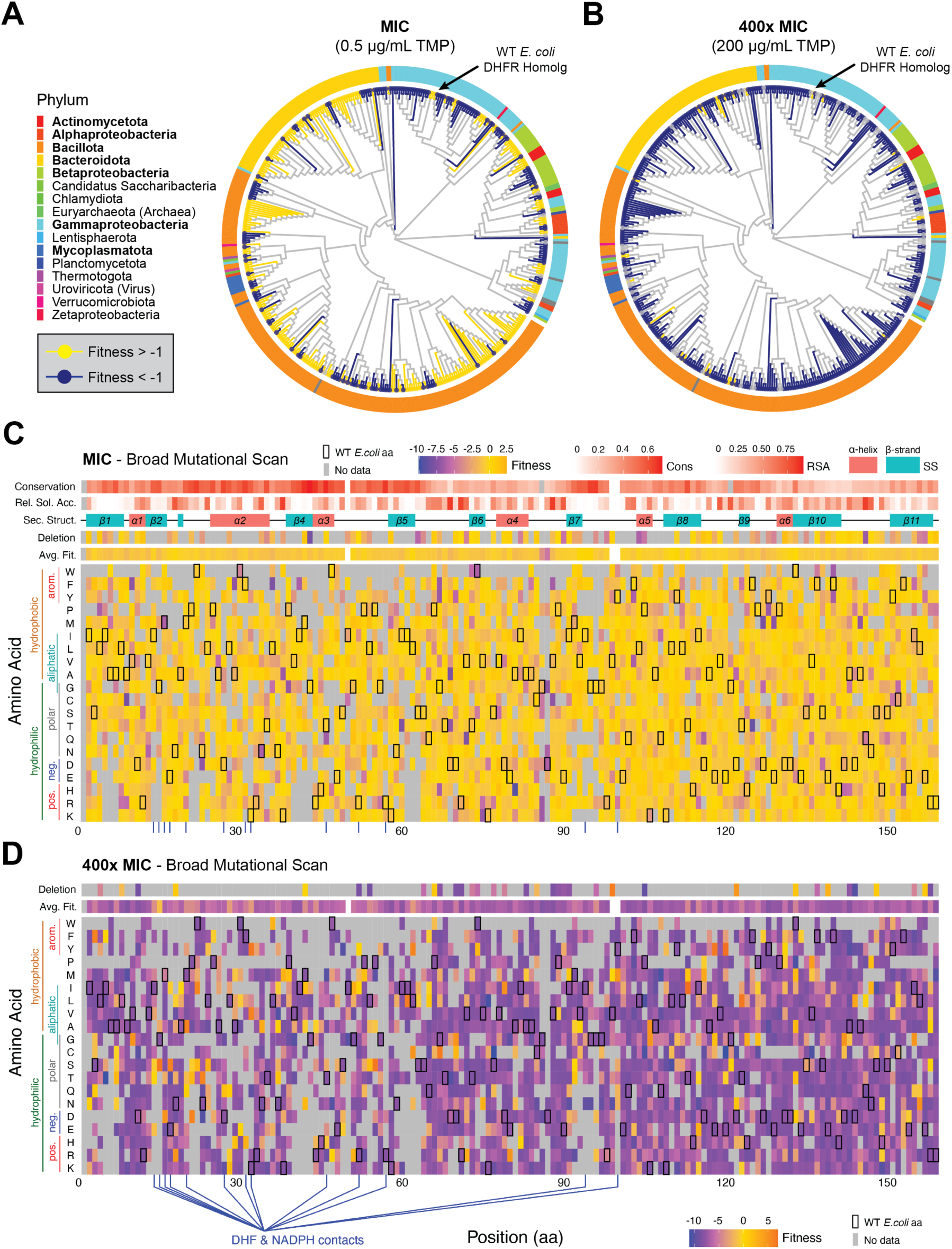
Trimethoprim Fitness: Phylogenetic Distribution and Broad Mutational Scanning Analysis. (**A**) Maximum likelihood phylogenetic tree of 416 DHFR homologs, showing trimethoprim-resistant homologs (fitness > −1) at minimal inhibitory concentration (MIC, 0.5 µg/mL TMP) in yellow, and non-resistant homologs (fitness < −1) in blue. Some degree of clustering can be observed. (**B**) Maximum likelihood phylogenetic tree of 416 DHFR homologs at 400× MIC (200 ug/mL TMP), with resistant homologs in yellow and non-resistant in blue. (**C**) Fitness landscape of 416 homologs and their associated mutants (within a 5-amino-acid range) mapped onto the wild-type *E. coli* DHFR sequence at MIC, with each homolog initially capable of complementing function in non-supplemented media. (**D**) Fitness landscape of 416 homologs and their associated mutants (within a 5-amino-acid distance) mapped onto the wild-type *E. coli* DHFR sequence at 400× MIC. Both heat maps show the average fitness of an amino acid substitution at each position along the protein sequence, with WT residues marked by black squares, and data on average fitness, deletions, secondary structure, relative solvent accessibility, and conservation included. Gray squares indicate no amino acid data.

At the high TMP concentration (200 µg/mL), only 27 homologs in the Codon 1 library were able to maintain function while resisting trimethoprim inhibition (fitness ≥ −1; **Table S6**). As with the MIC results, fitness differences between homolog pairs at this higher concentration varied significantly across the phylogenetic tree, influenced by evolutionary distance (**Fig. 5B**). Statistical support at the high TMP concentration was comparable to the MIC level, where the Wilcoxon rank-sum test still indicated a highly significant difference in fitness variation (W = 2.8 × 10⁸, *P* < 2.2 × 10⁻¹⁶), and Welch’s t-test confirmed a lower mean absolute fitness difference for closely related homologs compared to distantly related homologs (t = −26.2, df = 6,206, *P* < 2.2 × 10⁻¹⁶; **Table S7**). These findings reinforce the trend observed at the MIC, suggesting that evolutionary relatedness continues to influence fitness similarity, with closely related species showing more comparable fitness responses even at high trimethoprim concentrations.

For the BMS analysis across the TMP gradient, we selected the same 416 homologs and their mutants (within a five-amino-acid distance) that successfully complemented function in non-supplemented media (fitness ≥ −1; see **Fig. 3C** for reference). This selection criteria focuses our analysis on functionally relevant variants under antibiotic pressure and enables a direct comparison of median fitness changes for each amino acid at each position across the gradient. These fitness scores are then mapped onto the *E. coli* DHFR sequence as a reference, contextualizing the results within a well-characterized model organism (**Figs. 5C, 5D**). The resulting heat maps illustrate the fitness landscape within the inherited sequence space, reflecting the natural sequence diversity of the DHFR protein family as it has evolved across species. Each point on the heat map represents the average effect of a specific amino acid–position combination, aggregated across diverse sequence backgrounds, offering insights into how sequence variation influences fitness.

Under MIC selection, most amino acid substitutions at each position across the DHFR protein sequence support sufficient fitness (≥ −1) to maintain function (median fitness = −0.4; **Fig. 5C**). However, certain positions exhibit a median fitness well below the −1 threshold, indicating greater depletion and suggesting that mutations at these residues are less tolerated, leading to reduced survival rates. Notably, residue 85 (median fitness = −3.2; Welch’s t-test = 7.0, df = 52, *P* = 5.43 × 10⁻⁹) and residue 86 (median fitness = −6.1; t = 6.1, df = 15, *P* = 2.14 × 10⁻⁵) show significantly lower fitness compared to other positions (see **Fig. S8**). An examination of the multiple sequence alignment used to map the high-dimensional data onto the *E. coli* reference revealed that positions 85 and 86 fall within gapped regions for most homologs, except for *E. coli*. This underscores a limitation of alignment-based dimensionality reduction, where gaps in homologous sequences can obscure functional constraints and introduce artifacts into fitness estimates. At 400× MIC selection, most amino acid substitutions across the protein sequence did not support sufficient fitness (≥ −1) to maintain DHFR function, with a median fitness of −8.2 (**Fig. 5D**).

As trimethoprim concentrations increased across the experimental gradient, amino acid diversity at each DHFR position declined, reflecting the progressive elimination of low-fitness variants under increasing antibiotic pressure. Specifically, the average number of tolerated amino acids per position decreased from 17 at complementation to 16 at MIC and only 14 at 400× MIC. This trend highlights the selective pressure exerted by trimethoprim, which progressively narrows the range of functional substitutions that can maintain DHFR activity. For example, at residue 85, the number of tolerated amino acids decreased from 10 under complementation to 7 at 400× MIC. Similarly, residue 86 showed a sharper decline in tolerated substitutions, dropping from 7 to 3, indicating that previously functional substitutions become increasingly incompatible with DHFR activity under higher trimethoprim concentrations. This reduced tolerance highlights critical residues essential for the enzyme’s ability to withstand antibiotic stress, as evidenced by the loss of less-fit variants.

### Resistance-Conferring Mutations in Pathogenic DHFR Homologs

Our third objective was to identify specific mutations within pathogenic variants that contribute to antibiotic resistance. To achieve this, we designed our DropSynth-assembled DHFR library to include a broad range of obligate and opportunistic pathogens (see **Fig. 1B**), including most members of the ‘ESKAPE’ group [30]. These pathogens, including *Enterococcus faecium*, *Klebsiella pneumoniae*, *Acinetobacter baumannii*, and *Pseudomonas aeruginosa*, are responsible for a significant proportion of antibiotic-resistant infections in clinical settings [31]. Our library also includes clinically relevant pathogens such as *Anaerococcus tetradius*, *Bacillus cereus*, *Salmonella enterica*, and *Vibrio cholerae*, as well as others with varying levels of susceptibility to trimethoprim, including *Haemophilus influenzae*, *Staphylococcus aureus*, and *Streptococcus pneumoniae* [3]. With this diverse collection, we conducted the first unified assessment of resistance-conferring mutations in pathogenic variants across the DHFR protein family.

While our DHFR library includes at least one homolog from each of the pathogens listed above, some variants failed to complement function in the *E. coli* Δ*folA*Δ*thyA* cells, resulting in a lack of fitness data for those homologs. Nevertheless, we successfully recovered fitness data for eight well-characterized pathogenic DHFR homologs from the complementation assay, allowing us to demonstrate the effectiveness of our approach in identifying resistance-conferring mutations (see **Fig. S9**). Among these, we focused on two homologs, *Bacillus cereus* and *Streptococcus pneumoniae*, which exhibited numerous mutant variants with diverse fitness responses across the TMP gradient. This enabled us to identify amino acid substitutions associated with trimethoprim resistance, shedding light on the molecular mechanisms underlying resistance development.

For *Bacillus cereus*, numerous random mutant variants of the reference sequence were generated through our DropSynth assembly, revealing a range of fitness responses across the TMP gradient (**Fig. 6A**). Among these, only one mutant, featuring a valine-to-alanine substitution at position 71 (V71A), conferred resistance to trimethoprim at all concentrations (**Fig. 6B**). This mutant exhibited increasing fitness across the gradient, whereas the reference sequence’s fitness declined (**Fig. 6A**).

**Figure 6.**
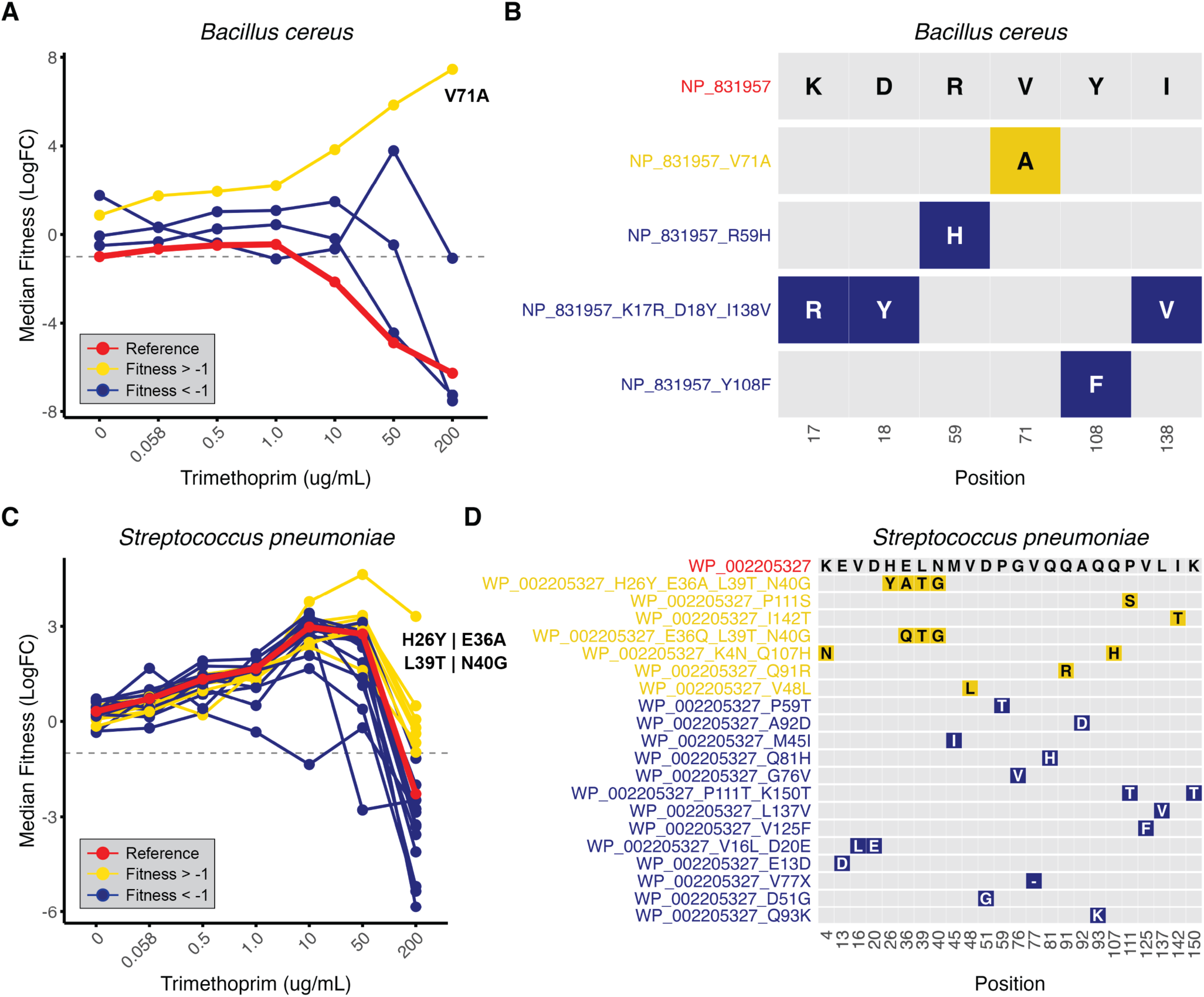
Resistance-Conferring Mutations in Pathogenic DHFR Homologs. (**A**) Median fitness of *Bacillus cereus* across the TMP gradient, showing the reference sequence in red, non-resistant mutants (fitness < −1) in blue, and TMP-resistant mutants in yellow. (**B**) Grid plot of amino acid substitutions in *B. cereus* (NP_831957; top row) showing unique mutant variants with single-letter amino acid codes and color scheme for fitness at high TMP (yellow for fitness ≥ −1, blue for fitness < −1). (**C**) Median fitness of *Streptococcus pneumoniae* across the TMP gradient, with the reference sequence in red, non-resistant mutants in blue, and TMP-resistant mutants in yellow. (**D**) Grid plot of amino acid substitutions in *S. pneumoniae* (WP_002205327; top row), with each row showing a unique mutant variant and colors indicating fitness at high TMP (yellow for fitness ≥ −1, blue for fitness < −1).

For *Streptococcus pneumoniae*, we identified seven unique mutant variants conferring resistance at high TMP concentrations (**Fig. 6C**). These included four single-substitution variants (V48L, Q91R, P111S, I142T), one double-substitution variant (K4N_Q107H), one triple-substitution variant (E36Q_L39T_N40G), and one quadruple-substitution variant (H26Y_E36A_L39T_N40G) (**Fig. 6D**). The triple- and quadruple-substitution variants shared mutations at residues 36, 39, and 40, but the quadruple mutant variant (H26Y_E36A_L39T_N40G) showed higher fitness across the TMP gradient (**Fig. 6C**). The remaining 13 mutant variants failed to restore metabolic function, indicating that substitutions, including a deletion (V77X), at these positions do not confer TMP resistance (**Fig. 6D**).

Remarkably, we recovered a perfect assembly *E. coli* DHFR homolog with a single amino acid substitution, lysine (K) to asparagine (N) at residue 38, which conferred increased resistance to trimethoprim relative to the WT *E. coli* reference homolog. This K38N variant showed improved resistance at higher trimethoprim concentrations, with fitness values greater than −1 at 50 µg/mL and 200 µg/mL (400× MIC), significantly outperforming the WT reference (see **Fig. S10**). As a perfect assembly homolog, we recovered 1,085 total barcodes for the K38N variant, including 169 high-quality barcodes with at least 10 sequences per barcode, ensuring reliable fitness data. These novel results reveal the robust fitness of the K38N variant across the TMP gradient, with dial-out PCR validation underway to confirm its role in conferring resistance at high TMP concentrations.

We also analyzed fitness data for homologs from the most abundant phyla in our DropSynth-assembled library, including Bacillota, Bacteroidota, and Alphaproteobacteria, Betaproteobacteria, and Gammaproteobacteria classes within Pseudomonadota (**Table S1**). Across 224 species and their mutant variants, we identified 40 species with mutations conferring resistance to high trimethoprim (TMP) concentrations (200 µg/mL). Specifically, resistance was observed in 1 of 6 Alphaproteobacteria species, 1 of 17 Betaproteobacteria species, 7 of 58 Gammaproteobacteria species, 18 of 92 Bacillota species, and 13 of 51 Bacteroidota species (**Figs. S11–S15**). This diverse spread of resistant mutations underscores the adaptive capacity and evolutionary plasticity of the DHFR enzyme, reflecting its divergent response to antibiotic pressure. Ultimately, these findings provide a comprehensive map of trimethoprim resistance across the DHFR protein family, revealing widespread antibiotic resistance and key genetic alterations across the bacterial domain.

### Multiplexed Fitness Validation Using Dial-out PCR

Our final objective was to validate pooled assay results using dial-out PCR [28] to measure the growth rates of individual homologs across the trimethoprim gradient. This validation step was necessary to confirm the accuracy of our high-throughput pooled assays and to ensure that the observed fitness effects were not artifacts of the competitive growth environment. Each DropSynth-assembled DHFR homolog is tagged with a unique barcode, enabling retrieval via PCR amplification. This barcoding system allows for precise identification and isolation of specific homologs from the pooled library, facilitating individual growth rate measurements (see **Methods**). We strategically targeted 10 DHFR homologs spanning a broad fitness range at 200 µg/mL TMP (median log₂ fold-change: −9.1 to +6.2; **Table S8**), ensuring representation of fitness profiles observed in the pooled assays. This selection included homologs from the pathogenic bacteria *Bacillus cereus* and *Streptococcus pneumoniae* (**Table S9**), which were of particular interest due to their clinical relevance and potential trimethoprim resistance (**Fig. 6**). WT *E. coli* DHFR and mCherry served as positive and negative controls, respectively. Growth rates of dial-out homologs were compared to their fitness values from pooled assays across the TMP gradient (**Fig. S16**).

As an example, the pathogenic homologs from *B. cereus* and *S. pneumoniae* exhibited growth rate changes across the TMP gradient that corresponded closely to their fitness responses observed in the pooled assays (**Figs. 7A, 7B**). These pathogenic dial-out variants, while distinct from the reference variants in the resistance analysis (**Fig. 6**), followed similar fitness trends across the TMP gradient and were suitable for dial-out validation (see **Fig. S17**). Specifically, the relatively stable growth rates for these homologs across the TMP gradient aligned with their consistent fitness responses to increasing trimethoprim concentrations, underscoring their inherent resistance to the antibiotic at low- to mid-concentrations (fitness ≥ −1 up to 50 µg/mL TMP). In contrast, the WT *E. coli* DHFR homolog exhibited declining growth rates across the gradient, corresponding to its progressively decreasing fitness response in the pooled assays due to trimethoprim inhibition (**Fig. 7C**). Spearman’s correlation analysis of all dial-out homologs demonstrated significant positive relationships between growth rates and fitness across the TMP gradient (e.g., rₛ = 0.67, *P* = 0.028 at 0 µg/mL TMP; rₛ = 0.75, *P* = 0.012 at 0.5 µg/mL TMP; rₛ = 0.67, *P* = 0.024 at 50 µg/mL TMP) (**Fig. S18**). These significant positive correlations between dial-out growth rates and corresponding fitness across TMP conditions provide independent validation for the multiplexed assay results, confirming the reliability of the pooled fitness measurements.

**Figure 7.**
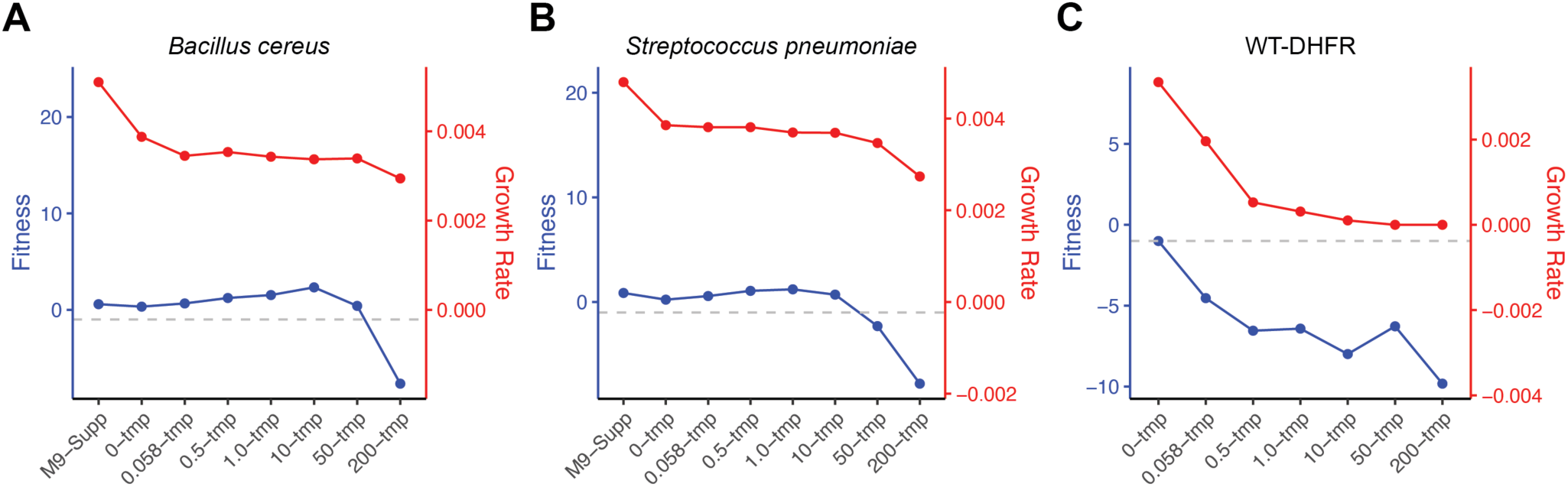
Dial-out Validation for Pathogenic DHFR Homologs. Line plots showing experimentally determined fitness from pooled assays (blue) and growth rates of individual homologs measured via dial-out PCR and plate reader assays (red) for the pathogenic homologs (**A**) *Bacillus cereus* and (**B**) *Streptococcus pneumoniae*, as well as the positive control (**C**) wild-type (WT) *E. coli* DHFR homolog. Growth rates (min⁻¹) were calculated as the maximum slope of OD₆₀₀ versus time on a log-linear plot.

## Discussion

This study presents the most comprehensive analysis of homolog complementation and inhibitor tolerance within the dihydrofolate reductase (DHFR) protein family to date. Using DropSynth technology [25,26], we assembled 1,208 unique DHFR homologs from two codon versions of a 1,536-member library, significantly advancing the scale of protein family analysis (**Fig. 1**). By combining this expansive library with multiplexed functional assays, we investigated DHFR responses to trimethoprim inhibition across a wide evolutionary range. This approach provides a more nuanced understanding of fitness landscapes and antibiotic resistance mechanisms than traditional Deep Mutational Scanning (DMS) [14,15,16], offering a broader and more holistic view of DHFR function and evolution. Our study enhances understanding of the evolutionary dynamics and functional diversity of the DHFR protein family, with implications for predicting resistance mechanisms and developing novel antibiotic strategies.

### Insights into Sequence-Function Relationships in DHFR Protein Evolution

To address our first objective, we conducted a complementation assay to evaluate the functionality of our assembled DHFR libraries in an *E. coli* knockout strain. The results demonstrated remarkable functional conservation of DHFR across diverse species, despite significant sequence divergence. Notably, more than half of all perfect assembly homologs successfully rescued the *E. coli* Δ*folA* phenotype, underscoring the enzyme’s evolutionary resilience (**Fig. 2B**). Furthermore, when we incorporated mutant variants with up to 5 amino acid changes, the rescue rate increased to over 90% (**Fig. 2C**). This substantial improvement in complementation efficiency highlights the adaptability of DHFR function through minor amino acid alterations, revealing the protein’s capacity to maintain its essential role despite sequence variations. However, we observed variations in complementation efficiency among codon-specific homologs, suggesting that even subtle sequence differences can significantly impact protein functionality (**Fig. 2D**). These findings underscore the complex interplay between sequence and function in the DHFR protein family and highlight the importance of considering both amino acid composition and codon usage when studying protein evolution and functionality in heterologous expression systems [41,42].

To investigate these effects, we used the RBS Calculator (version 2.1) [43] to predict the translation initiation rate for 364 homologs but found no clear trends, suggesting the involvement of more complex factors. DHFR is known to operate within a “Goldilocks zone” of protein expression, where both insufficient and excessive expression can adversely affect cell growth [33]. Numerous other effects likely combine to influence the measured growth rate beyond the catalytic activity of each homolog. These include the propensity of DHFR to adopt a molten globule state, which is influenced by the availability of chaperones (GroEL) and proteases (Lon) [29,44]. The coupling of DHFR to thymidylate synthase also plays a critical role, as mismatches in metabolic flux can lead to the accumulation of toxic intermediary products [36,45]. Finally, the modulation of protein-protein interactions has been proposed as another mechanism impacting cellular fitness [33]. These findings underscore the complex interplay between sequence, structure, and function in protein evolution, shedding light on how specific sequence changes influence functional outcomes.

Using Broad Mutational Scanning (BMS), an innovative approach for mapping the fitness landscape of protein homologs generated through large-scale gene synthesis, we gained key insights into the mutational landscape of DHFR homologs (**Fig. 3**). First, conserved sites exhibited significantly constrained mutational fitness, emphasizing their critical role in protein function, with a strong negative correlation (ρ = −0.51) indicating that mutations at these positions likely disrupt stability or function (**Fig. S3A**). Second, buried residues had smaller variability in fitness than solvent-accessible residues, possibly due to the potential impact of mutating buried residues on structure (**Fig. S3B**). Third, the correlation between higher mutational coverage and increased positional fitness suggests that highly deleterious mutations are less likely to be captured by the assay due to the loss of the variant or insufficient reads to pass quality filters (**Fig. S3C**). This effect may introduce a bias toward higher fitness values, as low-fitness homologs were excluded. Finally, the lack of significant differences in BMS fitness across secondary structures highlights the complexity of mutational effects, while also highlighting the potential variability in structural features among homologs (**Fig. S3D**). Projecting homologs onto the *E. coli* reference assumes structural conservation across the protein family, which may not fully account for such differences.

Our gain-of-function (GOF) analysis further revealed how specific mutations can rescue metabolic function under complementation conditions. For example, we identified nearly 500 GOF mutants with single amino acid substitutions associated with 191 dropout homologs (**Fig. 3A**). Mapping these mutations to the WT *E. coli* DHFR sequence (**Fig. 3E**) revealed key residues in the NADPH binding pocket, including positions 97 and 98, which form hydrogen bonds with the nicotinamide ring to aid catalytic positioning [46], and position 17 in the Met20 loop, which stabilizes substrate and cofactor binding [40,47]. GOF mutations included both buried and surface-exposed residues, underscoring the structural and functional importance of these sites in fitness recovery and their potential for evolutionary adaptation and therapeutic targeting.

### Complex Mechanisms Underlying Trimethoprim Resistance across DHFR Protein Family

For our second objective, the trimethoprim resistance assays revealed a spectrum of antibiotic susceptibility among DHFR variants, indicating that resistance operates along a continuum rather than as a binary trait (**Fig. 4**). Some variants exhibited moderate resistance, while others displayed high levels, suggesting that the degree of resistance is influenced by multiple factors, such as specific mutations in the DHFR gene, changes in enzyme activity, and alterations in substrate binding. These findings align with established mechanisms of TMP resistance, such as competitive binding of TMP to the folate pocket and the role of active-site substitutions in conferring resistance [11,48,49]. The decline in the number of resistant homologs with increasing TMP concentration reflects the progressive loss of DHFR function under TMP pressure (**Fig. 4D**), highlighting the complexity of resistance mechanisms as amino acid substitutions become less viable at higher drug levels (**Fig. 5D**). This spectrum challenges simplistic models of resistance and highlights the need for more sophisticated approaches in antibiotic development, such as targeting multiple variants or developing compounds that inhibit both wild-type and resistant forms [50,51].

The maintenance of functionality in a small subset of homologs at high TMP concentrations suggests the presence of resistance-conferring mutations. This aligns with previous findings that identify specific residue variations, such as D27E and L28Q, as common structural elements in TMP-resistant *E. coli* DHFR variants [11,52]. Furthermore, the fitness differences between closely related and distantly related homologs under TMP selection support the idea that resistance often emerges through specific mutations in the DHFR gene, with evolutionary proximity influencing the likelihood of sharing resistance-conferring mutations [53]. Here, the relationship between phylogenetic relatedness and fitness responses among DHFR homologs proved complex. While some closely related homologs exhibited similar fitness profiles (**Fig. 5A**), others showed unexpected divergence, especially as the selective pressure of TMP increased across the experimental gradient (**Fig. 5B**). This variability highlights the intricate nature of protein evolution and the challenges of predicting functional outcomes based solely on sequence similarity. These findings have significant implications for both clinical practice and evolutionary biology, emphasizing the importance of species-specific antibiotic selection in clinical settings and enhancing our understanding of resistance as an evolutionary adaptation.

### Identifying Key Resistance-Conferring Mutations in Pathogenic DHFR Variants

In our third objective, we identified mutations conferring trimethoprim resistance in pathogenic DHFR homologs, including members from the ESKAPE group [30,31]. This approach successfully highlighted key mutations in clinically relevant pathogens, such as *Bacillus cereus* (V71A) and *Streptococcus pneumoniae* (V48L, Q91R, P111S, I142T, and multi-substitution variants), demonstrating its power in uncovering the genetic basis of antibiotic resistance (**Fig. 6**). Notably, we discovered the K38N mutation in *E. coli* DHFR, which conferred enhanced resistance to trimethoprim at the highest concentrations tested (**Fig. S10**). Interestingly, a deep mutational scan of wild-type *E. coli* DHFR exposed to mild trimethoprim levels (3 µg/mL) also identified N38 variants, although at a much lower frequency than K38 (99.57% versus 0.14%) [54]. Our assay detected a smaller advantage in resistance for N38 at 1 µg/mL compared to higher TMP concentrations, suggesting a potential concentration-dependent effect of this substitution. This finding underscores the significant impact that even single amino acid substitutions can have on drug susceptibility, emphasizing the importance of monitoring minor genetic changes in bacterial populations, as they may contribute to the emergence of resistant strains. These results not only inform the design of new antibiotics or modifications to existing ones, potentially circumventing known resistance mechanisms, but also contribute to the development of improved diagnostic tools for rapid detection of resistant strains, enabling more targeted and effective treatment strategies.

### Validation of High-Throughput Fitness Measurements

For our final objective, we validated our pooled assay results by using dial-out PCR [28] to measure individual homolog growth rates across the trimethoprim gradient, providing critical support for the accuracy and reliability of our high-throughput approach. By strategically selecting 10 DHFR homologs that span a wide fitness range, including clinically relevant pathogenic variants, we were able to confirm that the fitness effects observed in the pooled assays were not artifacts of the competitive growth environment or compositional nature of the data. The similarity between individual growth rates and pooled fitness responses, particularly for pathogenic homologs from *B. cereus* and *S. pneumoniae* (**Fig. 7**), underscores the robustness of our experimental design. The significant positive correlations between dial-out growth rates and corresponding fitness across various TMP conditions, as revealed by Spearman’s correlation analysis (**Fig. S18**), provide strong independent validation for the multiplexed assay results. This not only confirms the reliability of our pooled fitness measurements but also strengthens the overall conclusions drawn from our study regarding the fitness landscape of DHFR homologs in response to trimethoprim stress.

### Power of DropSynth and Future Directions

Our approach, while similar to DMS analysis, focuses on naturally occurring homologs across diverse species rather than individual proteins, enabling broader exploration of evolutionary diversity and functional conservation within the DHFR family. By leveraging high-throughput, multiplexed gene synthesis and functional assays, we overcome traditional limitations in studying diverse protein families and uncover how these enzymes evolve, adapt, and develop antibiotic resistance. By examining a wide range of naturally occurring variants, we gain deeper insights into essential protein features, enhancing our understanding of DHFR’s catalytic mechanisms and substrate binding across species and physiological contexts, with significant implications for protein engineering, drug design, and molecular evolution.

Our findings also have significant implications for machine learning (ML) models in protein function prediction and generative models, particularly given that most current models rely solely on uncharacterized metagenomic sequences in an unsupervised manner [55,56]. Our observation that nearly half of all homologs failed to complement effectively highlights the importance of thorough functional characterization and underscores a well-documented challenge with the heterologous expression of proteins. Including non-complementing sequences in training datasets without adequate validation could introduce noise or bias, potentially poisoning the training set and compromising model accuracy [57,58]. This issue warrants further quantitative investigation, which will be a focus of our future work, to enhance the reliability of ML-based predictions in protein function and antibiotic resistance. Moreover, the observation that GOF mutations were present in nearly all non-complementing homologs offers a compelling opportunity for ML models to uncover the underlying patterns and mechanisms driving functionality.

Looking ahead, we aim to expand this approach to other essential bacterial enzymes and scale up to larger libraries, assembling over 10,000 homologs with millions of mutant variants simultaneously. Future work will also focus on developing more sophisticated modeling techniques to enhance the prediction of antibiotic resistance from sequence data and will be aided by the development of new analysis approaches less reliant on sequence-specific alignment. Additionally, investigating the structural basis of resistance mutations and designing narrow-spectrum antibiotics that selectively target pathogenic DHFR variants while preserving beneficial microbiota represent promising research directions. The utility of DropSynth and BMS demonstrated in this study opens new possibilities for exploring other protein families and resistance mechanisms, paving the way for more effective antibiotic development and personalized treatment strategies.

## Methods and Materials

### DropSynth Library Assembly

The methods for assembling the DropSynth libraries were previously described in detail by Sidore et al. [26]. Briefly, the process involves five major components: (1) Oligo Design, where a pool of 33,792 230-mer oligos was designed and synthesized, encoding two codon versions of a 1,536-member DHFR homolog library; (2) Barcoded Bead Design and Assembly, which involved creating unique 12-mer barcodes and assembling them onto magnetic beads; (3) Oligo Amplification and Barcoding, where oligo subpools were amplified, processed, and hybridized to the barcoded beads; (4) Emulsion PCR Assembly, during which the loaded beads were compartmentalized in droplets and underwent assembly via PCR; and (5) Bulk Suppression PCR, which involved amplifying the assembled gene libraries using specific primers and conditions to obtain the final product. See **Figure 1A** for a conceptual overview.

### pCVR205 Plasmid Construction

The DHFR expression vector for *E. coli* was derived from plasmid SMT205 (Addgene #134817), which supports leaky expression with lac-o repression, eliminating the need for IPTG induction to avoid toxic DHFR accumulation [29]. The plasmid was modified by removing unnecessary restriction sites originally used for cloning and reintroducing three sites compatible with the *folA* gene library (see **Fig. S19** for reference). NdeI and KpnI sites were removed from the wild-type *thyA* gene by leveraging genetic code degeneracy, and a gBlock was designed to insert one KpnI site and one BspQI site at the 3′ end of the DHFR gene. This gBlock was integrated into the SMT205 backbone using Golden Gate assembly, creating the plasmid pCVR1. The gBlock insertion was validated by Sanger sequencing using internal and external primers to the *thyA* region. Site-directed mutagenesis (SDM) was then used to introduce an NdeI site at the 5′ end of *folA*, resulting in the final construct, pCVR205. The plasmid was transformed into *E. coli* DH5α cells, and colonies were confirmed through Sanger sequencing. The final plasmid is available on Addgene (#198715). The gBlock sequence and primers used are provided in Supplemental Materials (**Table S10**).

### Ligation of Barcoded Libraries into pCVR205 Plasmid

DHFR libraries were ligated to the digested pCVR205 plasmid at a 3:1 insert-to-vector molar ratio, then purified and eluted. Approximately 170 ng of the ligated product from each library was transformed into 25 µL of DH5α cells, which were allowed to recover in SOB media before plating on LB agar with 10X dilutions to estimate colony numbers. The libraries were recovered by scraping the cells from the plates, pelleting them, and extracting plasmid DNA using the Monarch Plasmid DNA Miniprep Kit (New England Biolabs).

### Assembly Barcode Sequencing and Annotation

The assembly barcoded libraries were sequenced using five paired-end 600-cycle runs on an Illumina MiSeq. After PCR amplification with sequencing primers, amplicons were gel-extracted, quantified using an Agilent 2200 TapeStation, and pooled for sequencing with custom MiSeq primers. All primers are available in our previous study [26]. Individual FASTQ files were downsampled to 1,880,288 reads to address coverage biases, and adapter sequences were trimmed with bbduk [59]. Paired-end reads were merged using bbmerge [59], and a custom Python script generated consensus nucleotide sequences for each barcode, mapping them to their corresponding variants. Barcodes linked to multiple variants were filtered based on Levenshtein distance to identify contamination [60]. A consensus sequence was generated from the majority base call, and variants were imported into RStudio [61] for coverage and fidelity analysis. Assemblies represented refers to the number of assemblies corresponding to a perfect amino acid sequence, while percent perfect assemblies indicate the median percentage of perfect sequences from constructs with at least 100 assembly barcodes. Mutant homolog sequences were annotated by aligning the consensus nucleotide sequence against designed DHFR homologs, extracting the closest matches, and performing pairwise amino acid alignments to annotate mutations within five amino acids of the designed sequence.

### DHFR Library Transformation into Folate-Deficient E. coli Knockout Model

DHFR is an essential gene that converts 7,8-dihydrofolate (DHF) to 5,6,7,8-tetrahydrofolate (THF), a key step in deoxythymidine phosphate biosynthesis (see **Fig. S20**). For the complementation assay, barcoded DHFR libraries were transformed into the folate-deficient *E. coli* knockout strain *ER2566* Δ*folA*Δ*thyA* [32], which requires external folate and thymidine for growth. Controls included the wild-type (WT) *E. coli* DHFR variant, the low-function D27N variant [33], and mCherry, a non-catalytic red fluorescent protein [34]. Control plasmids were barcoded with 20 randomized nucleotides via In-Fusion HD cloning. Fifteen colonies from each control were sequenced and mixed with the library at a 0.1% molar ratio. The combined library and controls (50 ng) were transformed in quadruplicate into 80 µL of *ER2566* Δ*folA*Δ*thyA* cells by electroporation. After recovery in SOB media, cells were plated on LB agar with chloramphenicol and thymidine and grown at 37°C for 22 hours. The cells were then scraped, resuspended in supplemented M9 media (80 µM adenosine, 0.5 mM glycine, 4.5 mM inosine, 4.2 µM calcium pantothenate, 0.5 mM methionine, and 0.21 mM thymidine), diluted to OD600 of 3, and plated again on supplemented M9 media for 22 hours. Transformation efficiency was quantified using si× 10X serial dilutions plated on supplemented M9 media and incubated at 37°C for 22 hours.

### Complementation Assay

For the complementation assay, cells from the initial transformation on supplemented M9 media plates were scraped, washed via three centrifugation rounds at 2500 rcf for 5 minutes, and resuspended in minimal M9 media (M9 salts, 0.4% glucose, 2 mM magnesium sulfate, and 35 µg/mL chloramphenicol) to an OD600 of 3 units per mL. Stocks were prepared at a 1:1 ratio of cells to 50% glycerol and stored at −80°C. Five milliliters of the washed cells were plated on minimal M9 media and incubated at 37°C for 22 hours to evaluate the ability of DHFR variants to restore metabolic function in the *E. coli ER2566* Δ*folA*Δ*thyA* knockout strain without any supplementation. Resulting cells were scraped, resuspended in minimal M9 media, and miniprepped using a Monarch Plasmid DNA Miniprep Kit (New England Biolabs) for plasmid extraction and sequencing. This process was repeated for four rounds, with cells from each round re-plated to initiate the next. However, only first-round results were analyzed due to challenges in normalizing barcode recoveries across time points.

### Trimethoprim Selection Gradient

A trimethoprim gradient ranging from 0 to 400 times the minimum inhibitory concentration (MIC) of 0.5 µg/mL [62] was prepared in minimal M9 media, with concentrations of 0 µg/mL (complementation), 0.058 µg/mL, 0.5 µg/mL (MIC), 1.0 µg/mL, 10 µg/mL, 50 µg/mL, and 200 µg/mL (400× MIC). Washed and diluted cells (OD600 of 3 units per mL) were spread evenly on agar plates with the trimethoprim gradient and incubated at 37°C for 22 hours to evaluate the tolerance of DHFR variants to the antibiotic. Resulting cells were scraped, resuspended in minimal M9 media, and miniprepped using a Monarch Plasmid DNA Miniprep Kit (New England Biolabs) for plasmid extraction and sequencing. As with complementation, this process was repeated for four rounds of trimethoprim treatment, with cells from each round re-plated at the same trimethoprim concentration to initiate the next round. Only first-round results were analyzed due to challenges in normalizing barcode recoveries across time points.

### Barcode Sequence Screening

Plasmid barcodes recovered from all nine sampling conditions were independently amplified using unique primer sets: C1) LB media with chloramphenicol and thymidine, C2) minimal M9 media with supplements, C3) minimal M9 media without supplements (complementation), C4) 0.058 µg/mL trimethoprim, C5) 0.5 µg/mL trimethoprim (MIC), C6) 1 µg/mL trimethoprim, C7) 10 µg/mL trimethoprim, C8) 50 µg/mL trimethoprim, and C9) 200 µg/mL trimethoprim (400× MIC). PCR was performed to amplify plasmid barcodes using five primer pairs that introduced custom sequencing adapters and library indexes compatible with Illumina sequencing. Plasmid primer sets and sequencing primer pairs are provided in Supplemental Materials (**Table S11**). The resulting amplicons were size-selected via gel extraction, purified, pooled, and sequenced on an Illumina NovaSeq 6000 S4 (35-cycle run), generating 1,010,919,383 total sequence reads across both codon version libraries (**Table S2**). Barcodes for each sampling condition were clustered using Starcode [63] to collapse barcodes within a Levenshtein distance of 1 [60].

### Fitness Calculations

To reduce noise in calculating fitness changes between sampling conditions, we filtered barcodes to retain only those with at least 10 reads across the nine conditions (C1-C9). This filtering decreased the unique barcodes from 474,789 to 323,256, with 712,211,978 total reads successfully mapped to these barcodes across both codon version libraries (∼70% of total sequences) (**Table S2**). Fitness scores were calculated for each mapped sequence associated with at least one barcode. Read counts for each condition were normalized based on the total sequencing depth relative to condition C2 (growth on minimal M9 media with supplements). The log_2_ fold-change between each subsequent sampling condition (C3-C9) and condition C2 was then calculated for each barcode using the following equation:

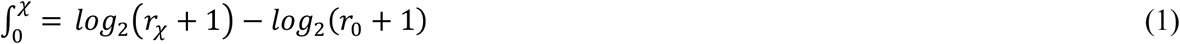

where *r*_χ_ is the number of normalized reads in the corresponding sampling condition (C3-C9) and *r*_0_ is the number of normalized reads in sampling condition C2. We then took the median value (to minimize effects of outliers) of the log_2_ fold-change for each unique DHFR variant based on its associated barcodes (BC) for each sampling condition using the following equation:

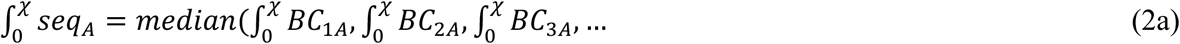

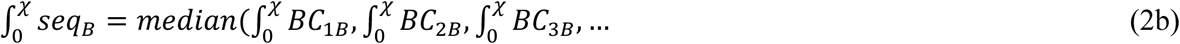

### Broad Mutational Scanning

Broad mutational scanning (BMS) was conducted on all homologs obtained from complementation (C3) and only those considered capable of complementation (median fitness > −1 at C3) across all trimethoprim selection conditions (C4-C9). We aligned the homologs using MAFFT and created a lookup table for each residue. For each perfect assembly homolog sequence (mutations = 0), we scanned all residues and recorded their fitness in a BMS data table, linking each residue to its corresponding *E. coli* position based on the alignment. For mutant variants within 5 amino acids of the designed homolog, we documented only the mutated residues and added their fitness to the BMS table. The BMS fitness for each residue was calculated as the median of all corresponding data points at that position.

### Gain-of-Function Mutations

To identify gain-of-function (GOF) mutations in the complementation assay, we classified DHFR homologs with fitness scores below −1 as dropout homologs and selected them for further analysis. These dropout homologs were screened for single amino acid mutant variants that restored function (fitness > −1), designating these as GOF mutants. We aligned the sequences of the GOF mutants to the WT *E. coli* DHFR sequence, counted the number of mutations at each position, and identified positions as significant if the mutation count exceeded two standard deviations above the mean. These significant GOF positions were mapped onto the *E. coli* DHFR protein structure (PDB 4KJK) [39] to visualize their spatial and functional relevance. Finally, we examined specific types of amino acid changes, including deletions, that contributed to fitness restoration at each significant position.

### DHFR Fitness Validation by Dial-out PCR

Each DHFR assembly is tagged with a unique barcode, enabling the retrieval of specific homologs from each codon version library through PCR amplification, referred to as dial-out PCR [28]. The dial-out procedure aims to independently verify an individual homolog’s fitness response to trimethoprim in isolation, without competition effects from the pooled multiplex assay. The WT *E. coli* DHFR gene (NP_414590) served as a positive control, while mCherry was our negative control. Dial-out PCR primers were designed to flank each homolog construct, with reverse primers annealing to the gene-specific barcode (**Table S12**). After gel extraction, individual amplicons were subjected to restriction digest using KpnI-HF and NdeI, ligated into an empty pCVR205 plasmid, transformed into electrocompetent *E. coli ER2566* Δ*folA*Δ*thyA* cells, and incubated in M9 minimal media with or without trimethoprim on 96-well plates. OD600 measurements were recorded every 6 minutes for 22 hours using a Synergy Neo2 microplate reader (Agilent BioTek) to calculate growth rates.

### Statistical Analysis

Data analysis and statistical tests, including Pearson’s (ρ) and Spearman’s (r_s_) correlations, Wilcoxon rank-sum tests, and Welch’s t-tests, were performed in R (version 4.3.2) [61]. Data visualizations were generated using the ggplot2 [64] and ggtree [65] packages, while structural visualizations were created with UCSF Chimera [66]. Residue conservation was assessed via Jensen-Shannon divergence [67], with secondary structure and relative solvent accessibility derived from DSSP analysis [68,69] of the 1H1T structure [70]. Statistical significance was set at *P* ≤ 0.05.

## Acknowledgments

We thank Kim Reynolds, Samuel Thompson, Tanja Kortemme, and the GC3F Core (UO) for productive discussions. We thank Ben Burress for his assistance with protein structure modeling. Plasmid SMT205 was provided by Tanja Kortemme (Addgene #134817).

## Funding

National Institutes of Health grant DP2TR004215 (CP)

University of Oregon Knight Campus Undergraduate Scholar fellowship (CR)

## Author contributions

Conceptualization: CR, CP.

Formal analysis: KJR, CR, CP.

Investigation: KJR, CR, SRH, CP.

Methodology: KJR, CR, SRH, CP.

Visualization: KJR, CP.

Resources: CP.

Supervision: CP.

Writing-original draft: KJR, CR, CP.

Writing-review and editing: KJR, CR, SRH, CP.

Project administration: CP.

Validation: KJR, CP.

Funding acquisition: CP.

## Competing interests

The authors declare competing financial interests. CP and SRH are co-founders of SynPlexity, a startup company commercializing DropSynth gene synthesis technology (https://synplexity.com).

## Data and materials availability

Sequence reads for synthetic DHFR homologs were submitted to the NCBI Sequence Read Archive under BioProject accession PRJNA1189478 (https://www.ncbi.nlm.nih.gov/bioproject/1189478). Processed read counts and R Markdown (RMD) files containing all analysis scripts for full reproducibility are available at the Plesa Lab GitHub (https://github.com/PlesaLab/DHFR). This repository houses the comprehensive dataset and code used in the DHFR study, enabling other researchers to replicate the analyses and build upon the findings. The processed read counts provide the foundation for the statistical analyses, while the RMD files offer step-by-step documentation of the data processing, statistical methods, and visualization techniques employed in the study. Material requests should be directed to the corresponding author: Calin Plesa.

## Supplementary Materials

This PDF file includes Figs. S1 to S20 and Tables S1 to S12.

## Notes

https://github.com/PlesaLab/DHFR

http://dx.doi.org/10.6084/m9.figshare.28266890

https://www.ncbi.nlm.nih.gov/bioproject/1189478

https://dropsynth.org

### Competing Interest Statement

CP and SRH are co-founders and hold equity in SynPlexity.

