## Supplementary Information for "Exploring Antibiotic Resistance in Diverse Homologs of the Dihydrofolate Reductase Protein Family through Broad Mutational Scanning"

Romanowicz† and Resnick† *et al.*

† **First Authorship:** These authors contributed equally to this work and share first authorship

**This PDF file includes:**

Figs. S1 to S20
Tables S1 to S12

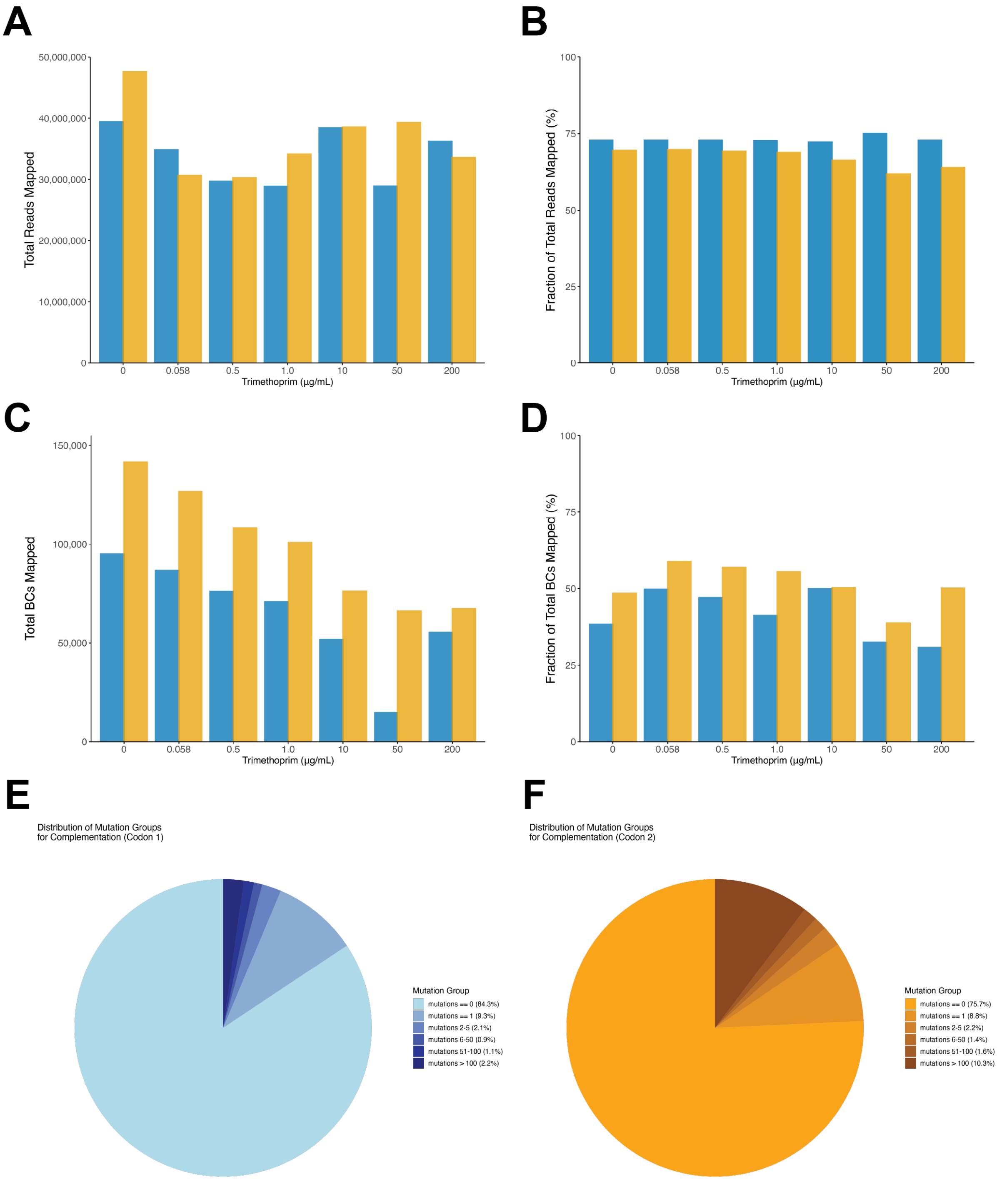

**Fig. S1. Sequencing Statistics Across TMP Gradient.** Sequencing reads and barcode statistics across the TMP gradient showing (**A**) total reads mapped at each TMP condition, (**B**) fraction of total reads mapped, (**C**) total barcodes (BCs) mapped, and (**D**) fraction of total BCs mapped. (**E**) The distribution of total reads mapped to barcodes by mutations in the Codon 1 library. (**F**) The distribution of total reads mapped to barcodes by mutations in the Codon 2 library.

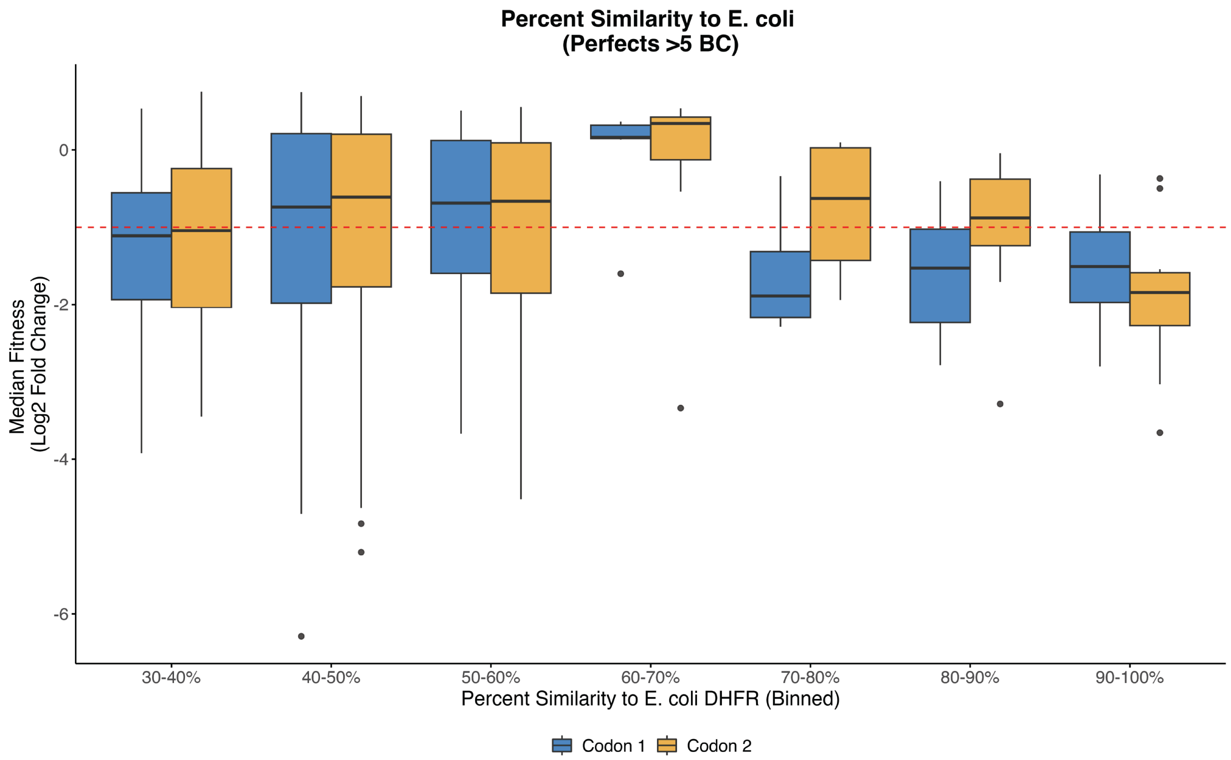

**Fig. S2. Sequence Similarity of DHFR Homologs to Wild-Type *E. coli* DHFR.** The median sequence identity of perfect assembly homologs recovered from the non-supplemented complementation condition was 46%. All DHFR homologs were grouped by percent similarity to the WT *E. coli* DHFR, with median fitness and standard deviation shown for each bin. Most distant homologs retained the ability to complement (fitness > -1) the function of native *E. coli* DHFR in the knockout strain. The red dashed line at -1 indicates the minimum fitness threshold for successful complementation.

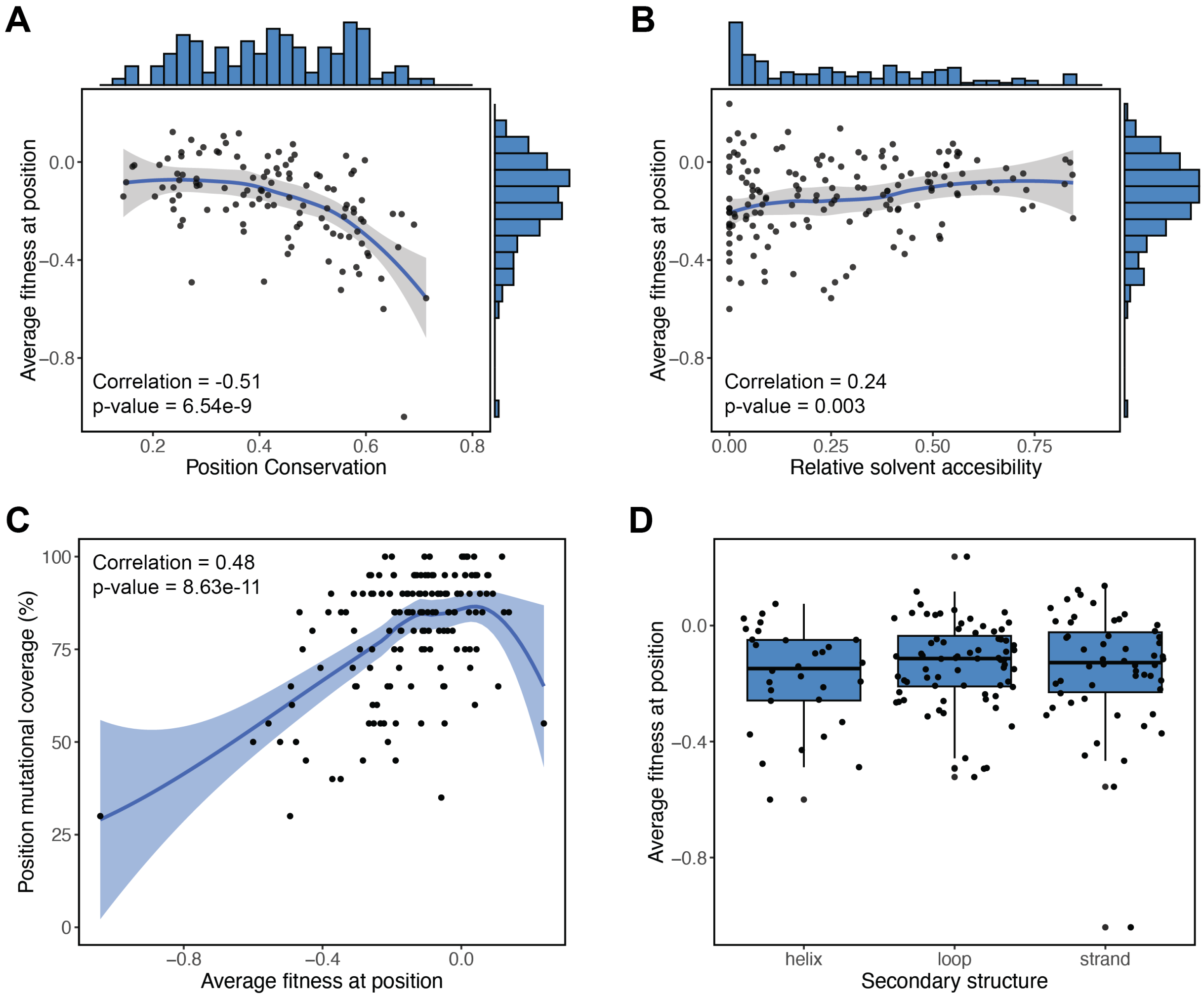

**Fig. S3. Broad Mutational Scanning Summary Statistics at Complementation.** (**A**) Average BMS position fitness compared to site conservation (Jensen-Shannon divergence) under complementation. (**B**) The average BMS position fitness compared to the relative solvent accessibility based on a DSSP analysis of the 1H1T crystal structure. Buried residues tend to be more constrained ($P = 0.003$). (**C**) Mutational scanning coverage decreases at sites of low fitness ($P = 8.63e^{-11}$) due to assembly barcodes with low read numbers. (**D**) Average BMS position fitness based on secondary structures in the E. *coli* DHFR structure (helix, loop, strand).

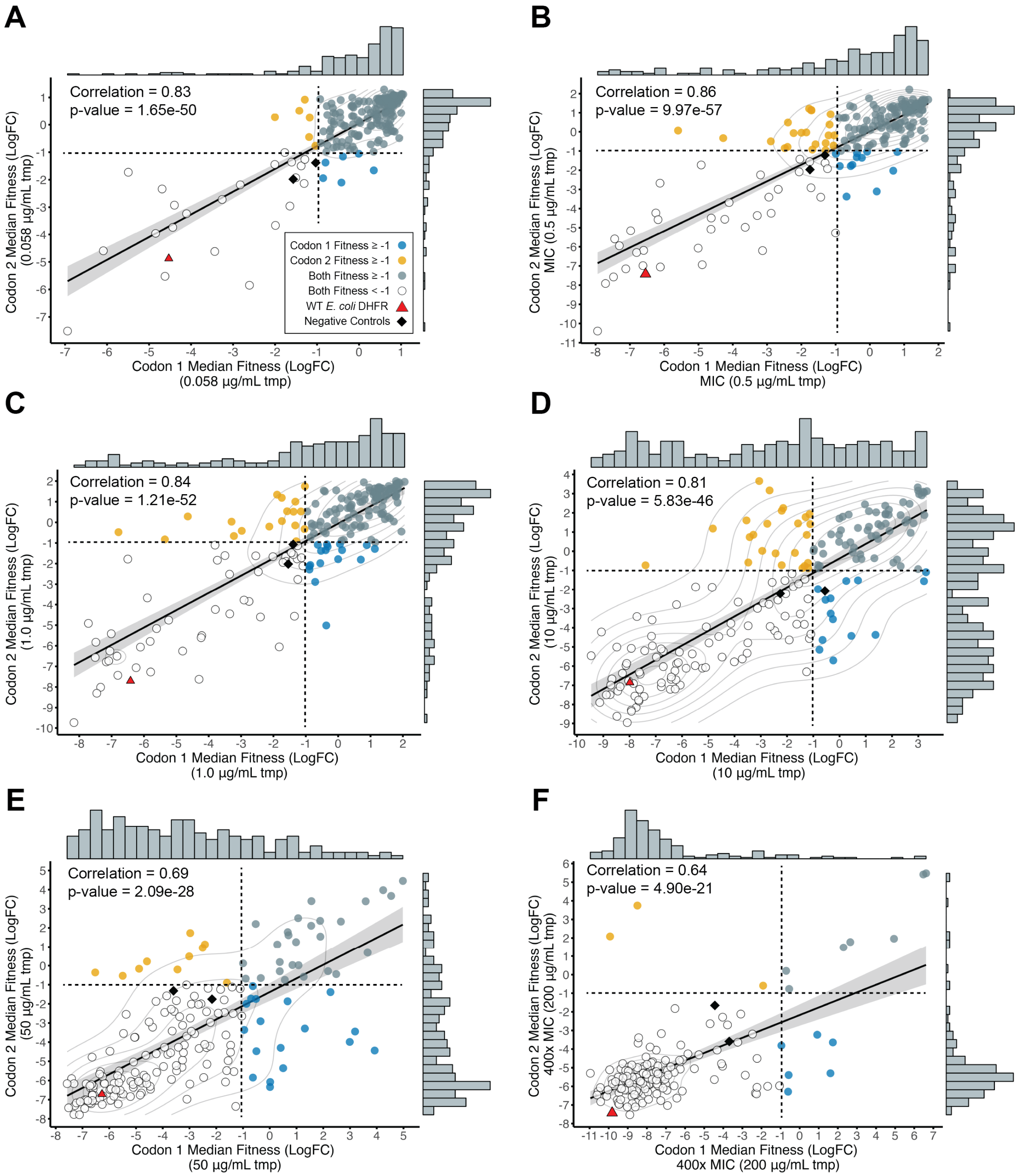

**Fig. S4. Fitness Correlations for Homologs Shared Between Codon Versions Along the TMP Gradient.** Pearson fitness correlations for shared homologs between codon versions at (**A**) 0.058 µg/mL TMP, (**B**) 0.5 µg/mL TMP (MIC), (**C**) 1.0 µg/mL TMP, (**D**) 10 µg/mL TMP, (**E**) 50 µg/mL TMP, and (**F**) 200 µg/mL TMP (400× MIC). Points are colored based on fitness for codon-specific homologs, shared homologs, or those with low fitness. The WT *E. coli* DHFR homolog and negative controls (D27N, mCherry) are plotted for reference. Dashed lines denote the minimum fitness thresholds for trimethoprim resistance.

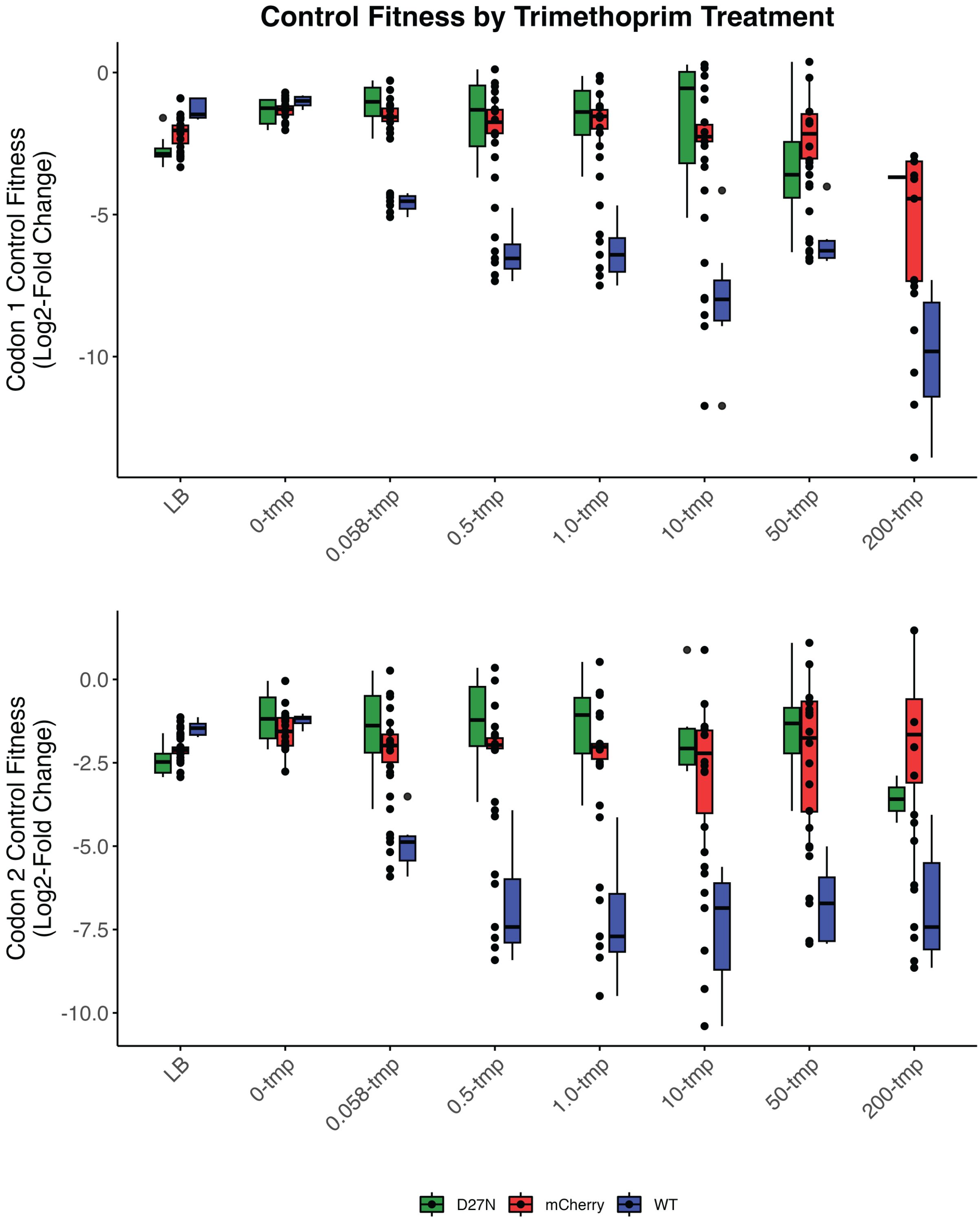

**Fig. S5. Control Response to Trimethoprim.** Boxplot illustrating homolog fitness across the trimethoprim gradient for Codon 1 (**top**) and Codon 2 (**bottom**), including the wild-type (WT) *E. coli* DHFR (positive control) and the negative controls, D27N and mCherry.

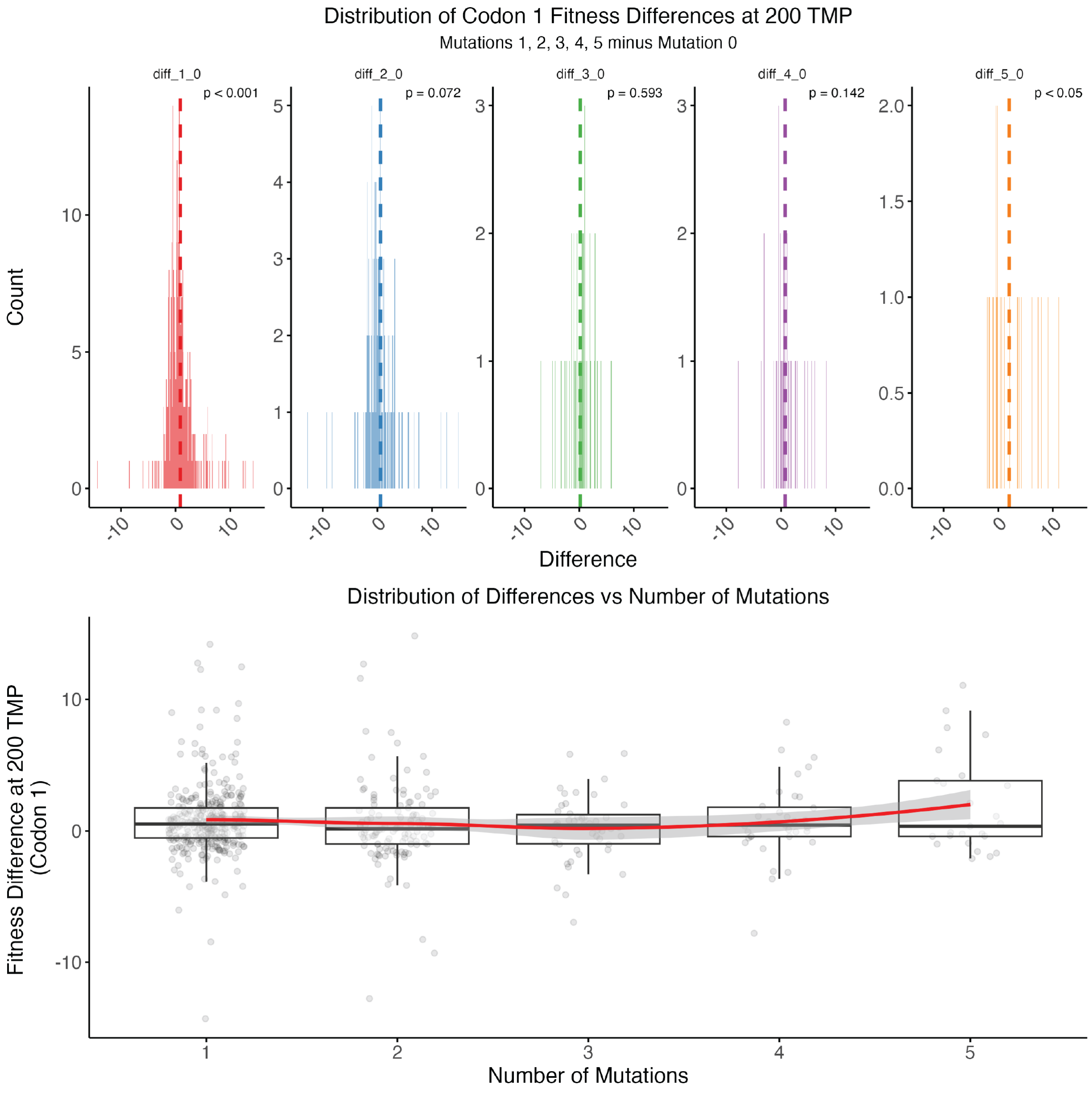

**Fig. S6. Codon 1 Fitness Differences by Mutations at 200 µg/mL Trimethoprim.** Distribution of mutant fitness responses at 200 µg/mL TMP for Codon 1, showing significant differences between single amino acid mutations and the perfect assemblies ($p < 0.001$), as well as for mutations with 5 amino acid differences compared to the perfect assemblies ($p < 0.05$). Mutants with 5 amino acid differences exhibit a significant positive impact on fitness at 200 µg/mL TMP.

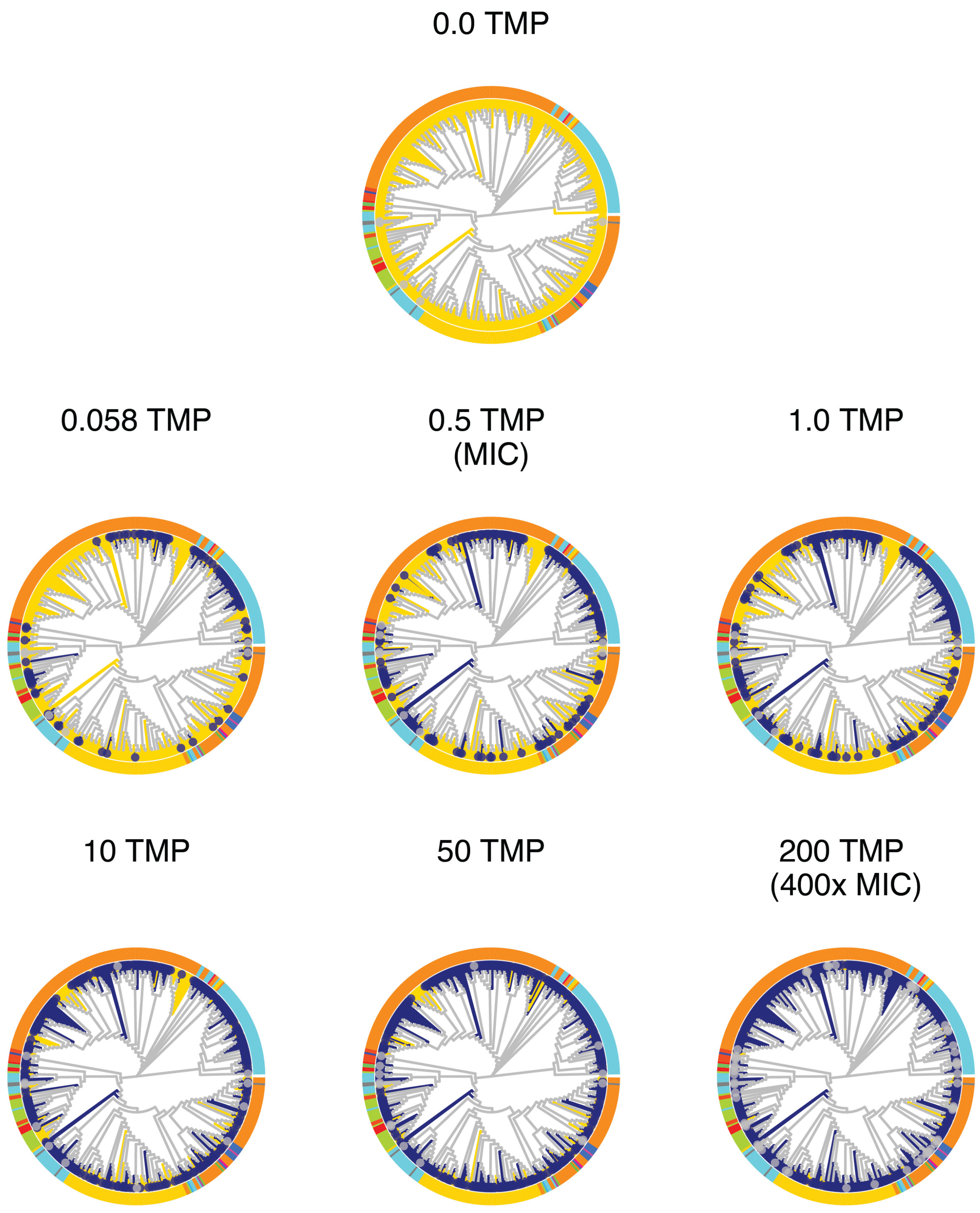

**Fig. S7. Phylogenetic Fitness Distribution of DHFR Homologs Across Trimethoprim Gradient.** Maximum likelihood phylogenetic trees of 416 DHFR homologs, showing trimethoprim-resistant homologs (fitness > -1) in yellow, and non-resistant homologs (fitness < -1) in blue across the trimethoprim (TMP) gradient.

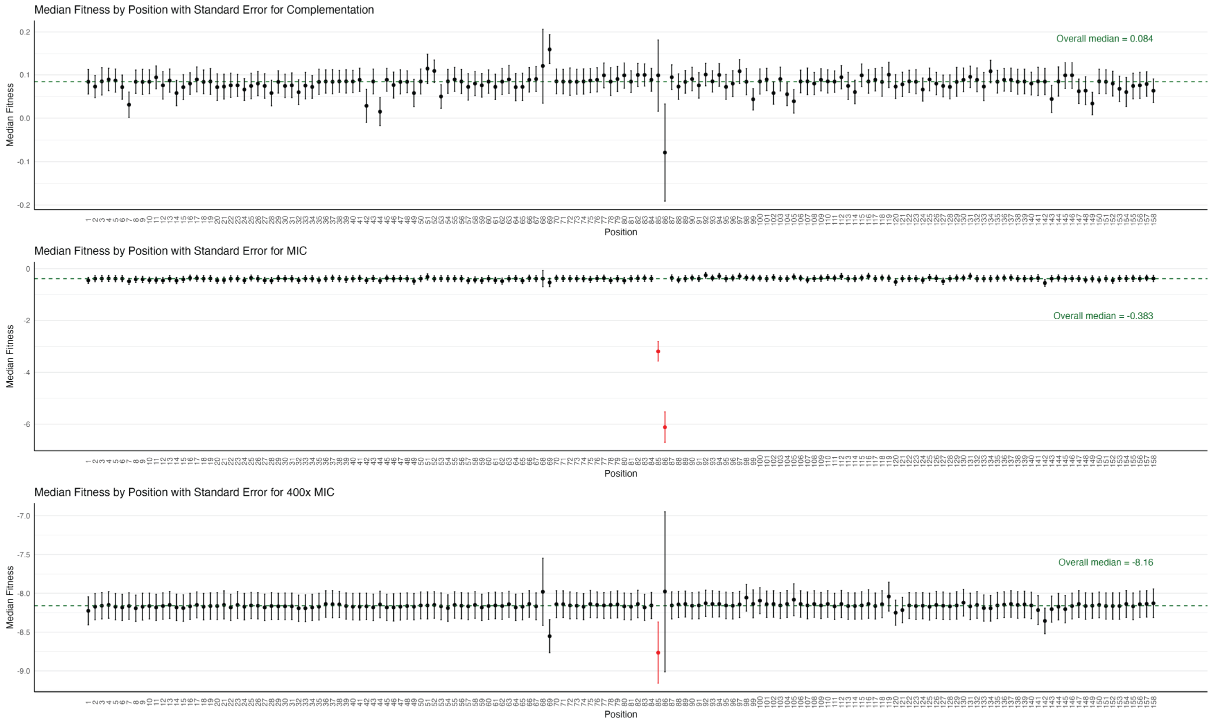

**Fig. S8. Median Fitness by Position Across TMP Gradient.** Points indicate the median fitness for recovered amino acids at each position under complementation (**top**), MIC (**middle**), and 400× MIC (**bottom**) conditions, with error bars representing the standard error of the median fitness. Red points highlight positions with median fitness significantly different from all others under those conditions. The green dashed line shows the overall median fitness across all positions.

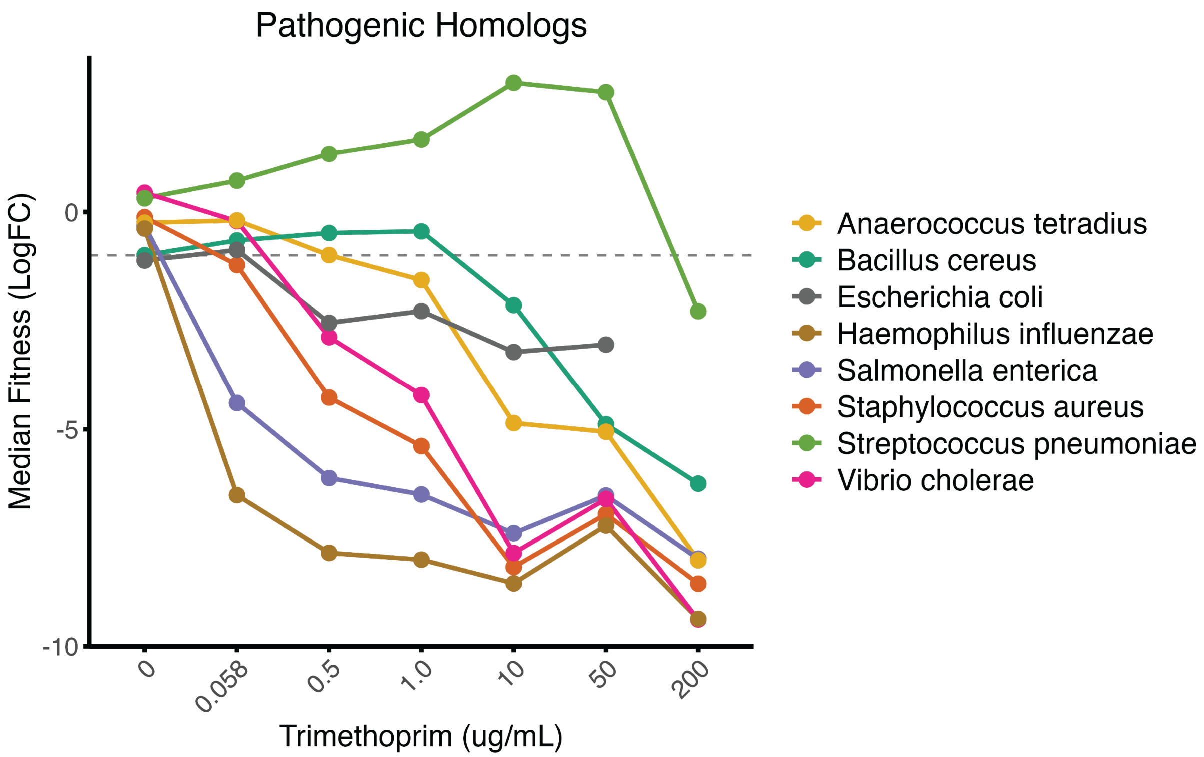

**Fig. S9. Trimethoprim Fitness for Pathogenic DHFR Homologs.** Median fitness for DHFR homologs across the trimethoprim gradient corresponding to the following pathogenic variants: *Anaerococcus tetradius*, *Bacillus cereus*, *Haemophilus influenzae*, *Salmonella enterica*, *Staphylococcus aureus*, *Streptococcus pneumoniae*, and *Vibrio cholerae*. The wild-type (WT) *E. coli* DHFR homolog is included as the positive control.

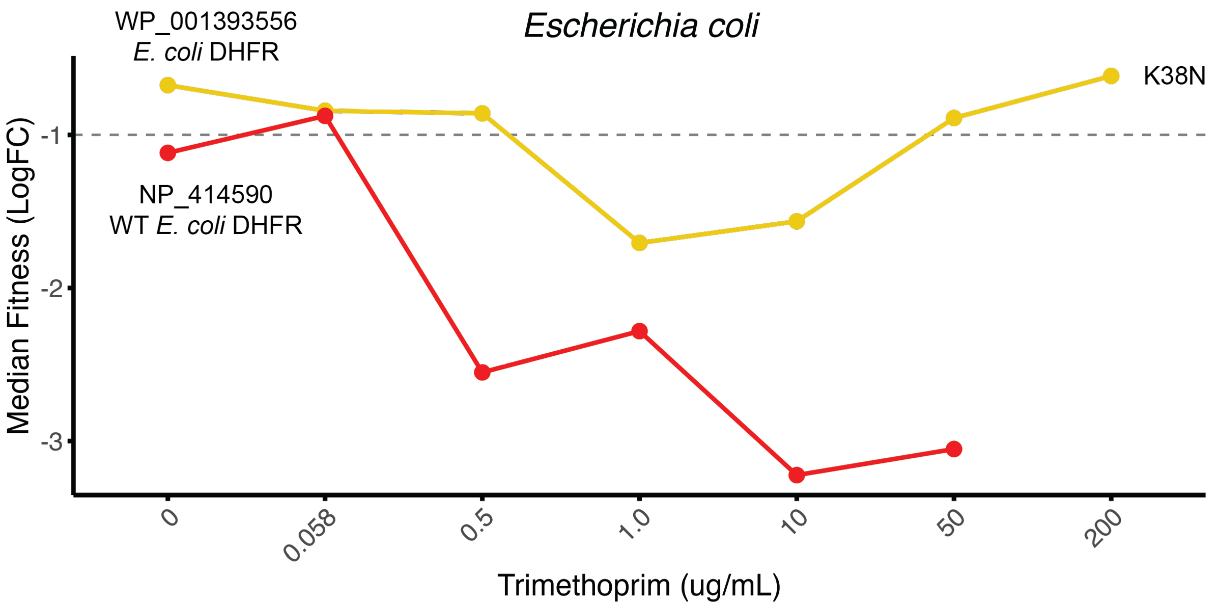

**Fig. S10. Fitness of K38N *E. coli* DHFR Homolog Relative to WT *E. coli* Homolog.** Comparison of median fitness between the perfect assembly *E. coli* DHFR variant (WP_001393556) and the wild-type (WT) *E. coli* DHFR variant (NP_414590) across the trimethoprim (TMP) concentration gradient. A single amino acid mutation (K38N) distinguishes the two variants, conferring trimethoprim resistance across the gradient relative to the WT reference.

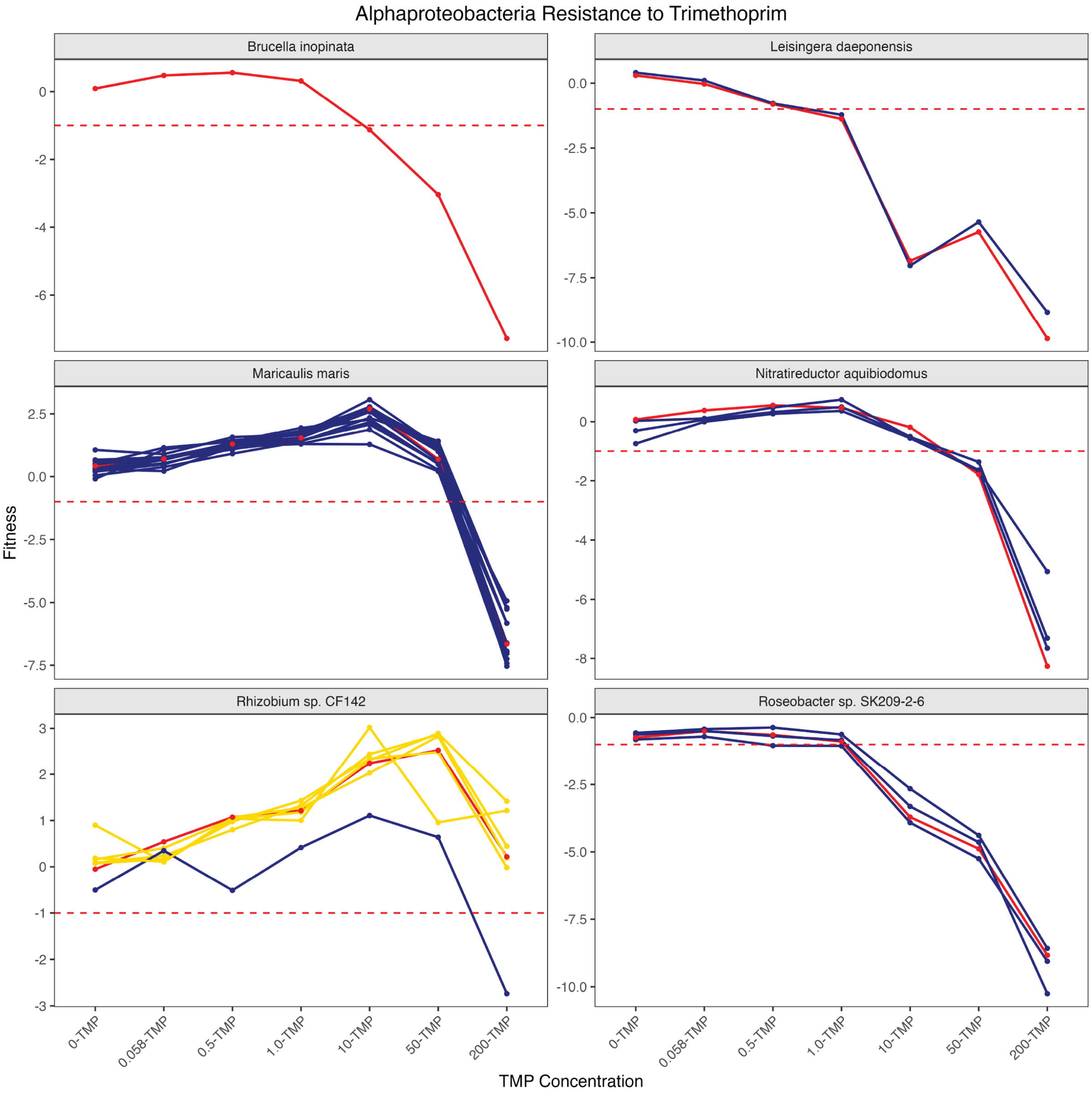

**Fig. S11. Alphaproteobacteria Fitness.** Median fitness (Log$_2$ fold-change) of six recovered Alphaproteobacteria species across the TMP gradient, showing the reference sequence in red, non-resistant mutants (fitness < -1) in blue, and TMP-resistant mutants (fitness > -1) in yellow. Only species with complete fitness data for the reference sequence and associated mutants were retained.

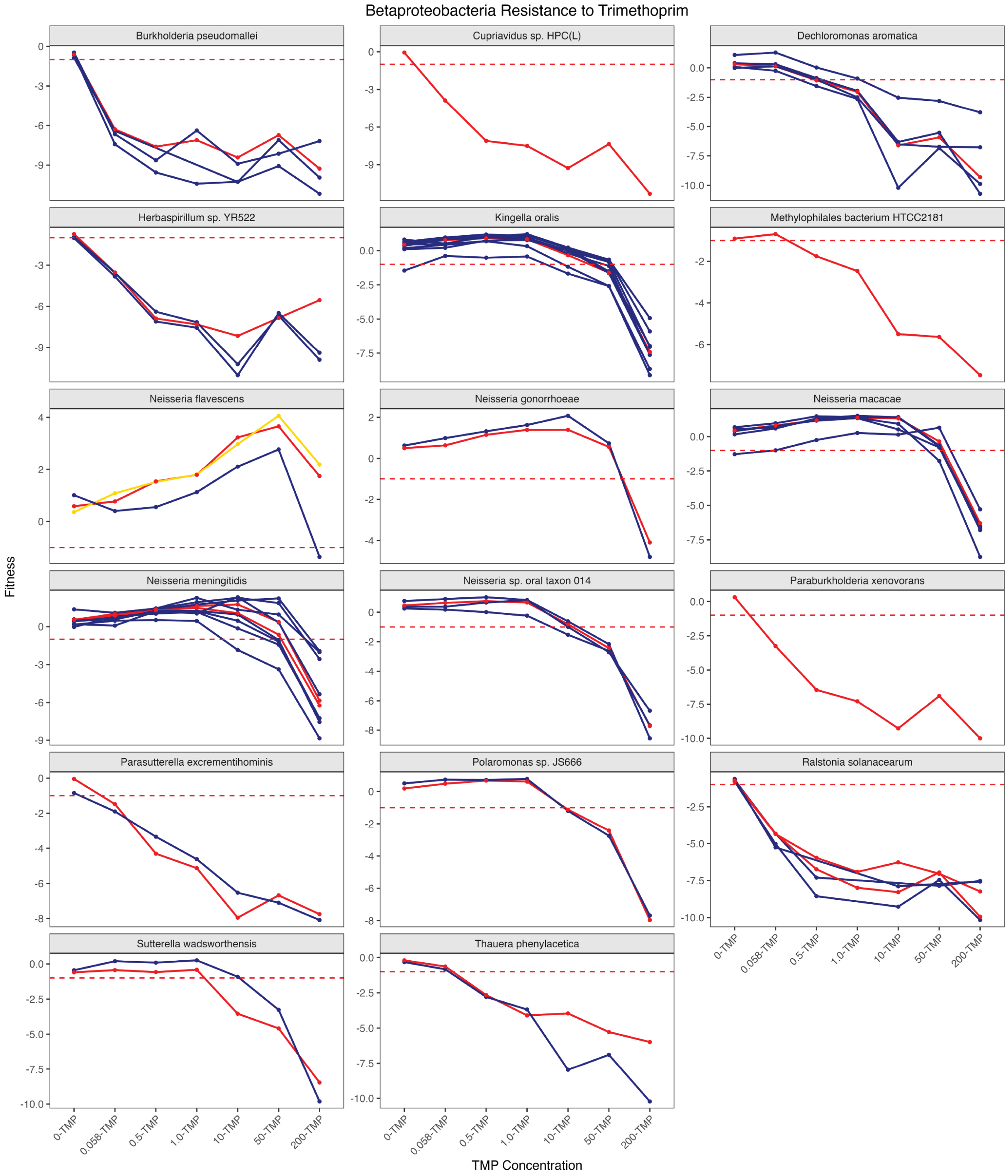

**Fig. S12. Betaproteobacteria Fitness.** Median fitness of 17 recovered Betaproteobacteria species across the TMP gradient, showing the reference sequence in red, non-resistant mutants (fitness < -1) in blue, and TMP-resistant mutants (fitness > -1) in yellow. Only species with complete fitness data for the reference sequence and associated mutants were retained.

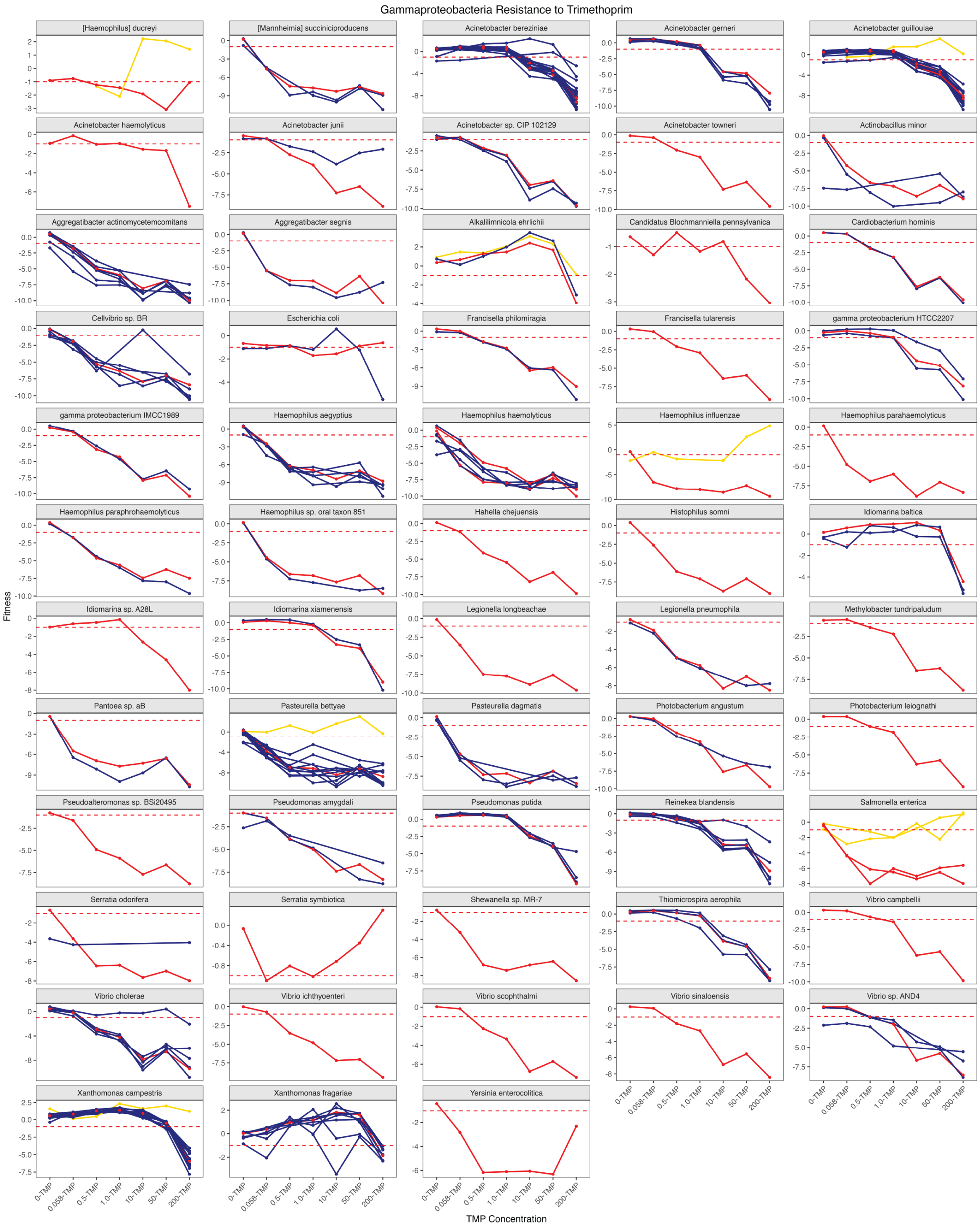

**Fig. S13. Gammaproteobacteria Fitness.** Median fitness of 58 recovered Gammaproteobacteria species across the TMP gradient, showing the reference sequence in red, non-resistant mutants (fitness < -1) in blue, and TMP-resistant mutants (fitness > -1) in yellow. Only species with complete fitness data for the reference sequence and associated mutants were retained.

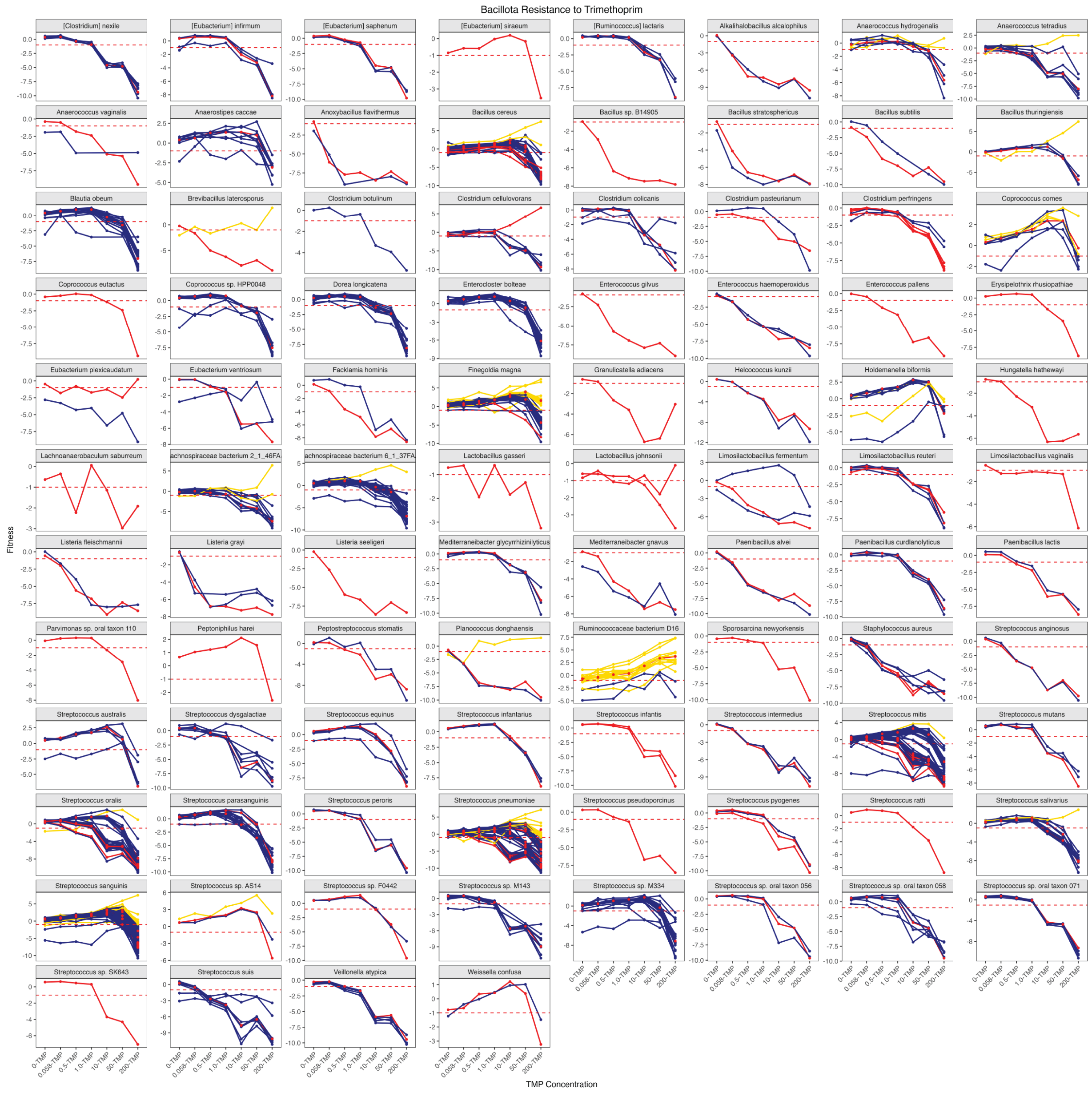

**Fig. S14. Bacillota Fitness.** Median fitness of 92 recovered Bacillota species across the TMP gradient, showing the reference sequence in red, non-resistant mutants (fitness < -1) in blue, and TMP-resistant mutants (fitness > -1) in yellow. Only species with complete fitness data for the reference sequence and associated mutants were retained.

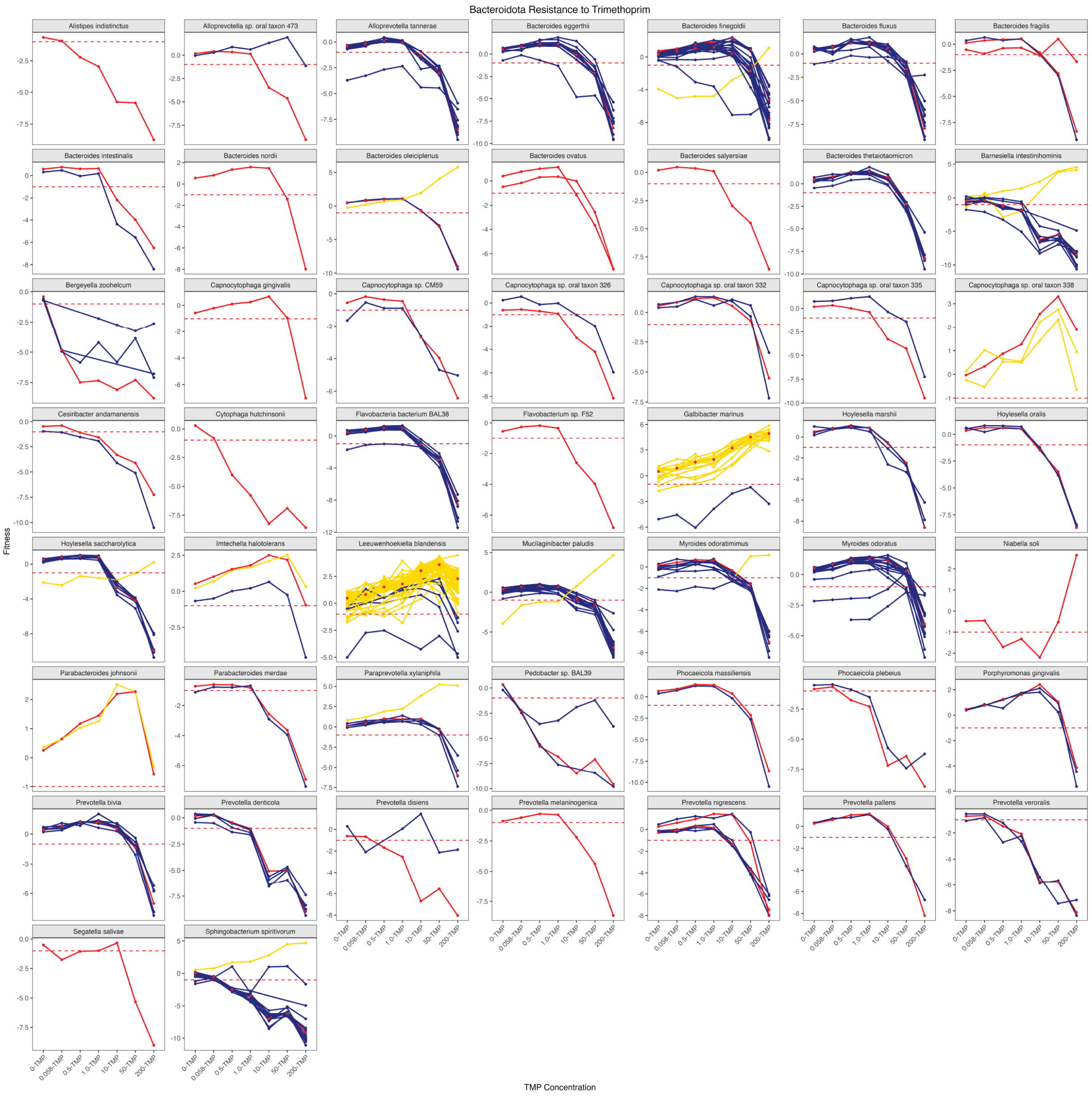

**Fig. S15. Bacteroidota Fitness.** Median fitness of 51 recovered Bacteroidota species across the TMP gradient, showing the reference sequence in red, non-resistant mutants (fitness < -1) in blue, and TMP-resistant mutants (fitness > -1) in yellow. Only species with complete fitness data for the reference sequence and associated mutants were retained.

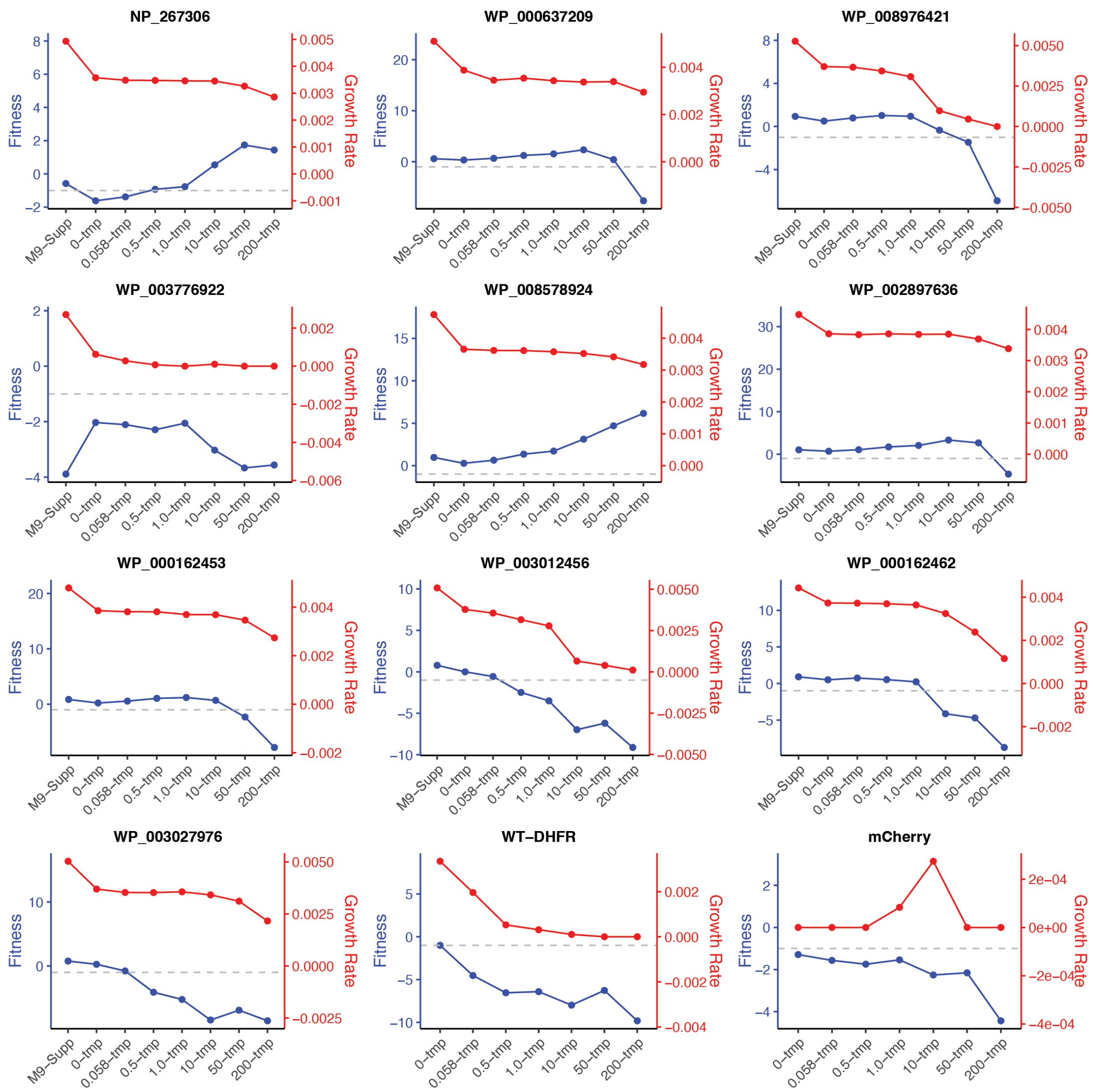

**Fig. S16. Comparison of Dial-out Fitness and Growth Rates.** Line plots showing experimentally determined fitness from pooled assays (blue) and growth rates of individual homologs measured via Dial-out PCR and plate reader assays (red). Growth rate ($min^{-1}$) is calculated as the maximum slope of OD600 vs. time on a log-linear plot. Plots represent 10 individual homologs spanning fitness values from strongly depleted (fitness < -9) to highly resistant (fitness > 6) under 200 µg/mL trimethoprim. Wild-type (WT) *E. coli* DHFR serves as the positive control, and mCherry as the negative control.

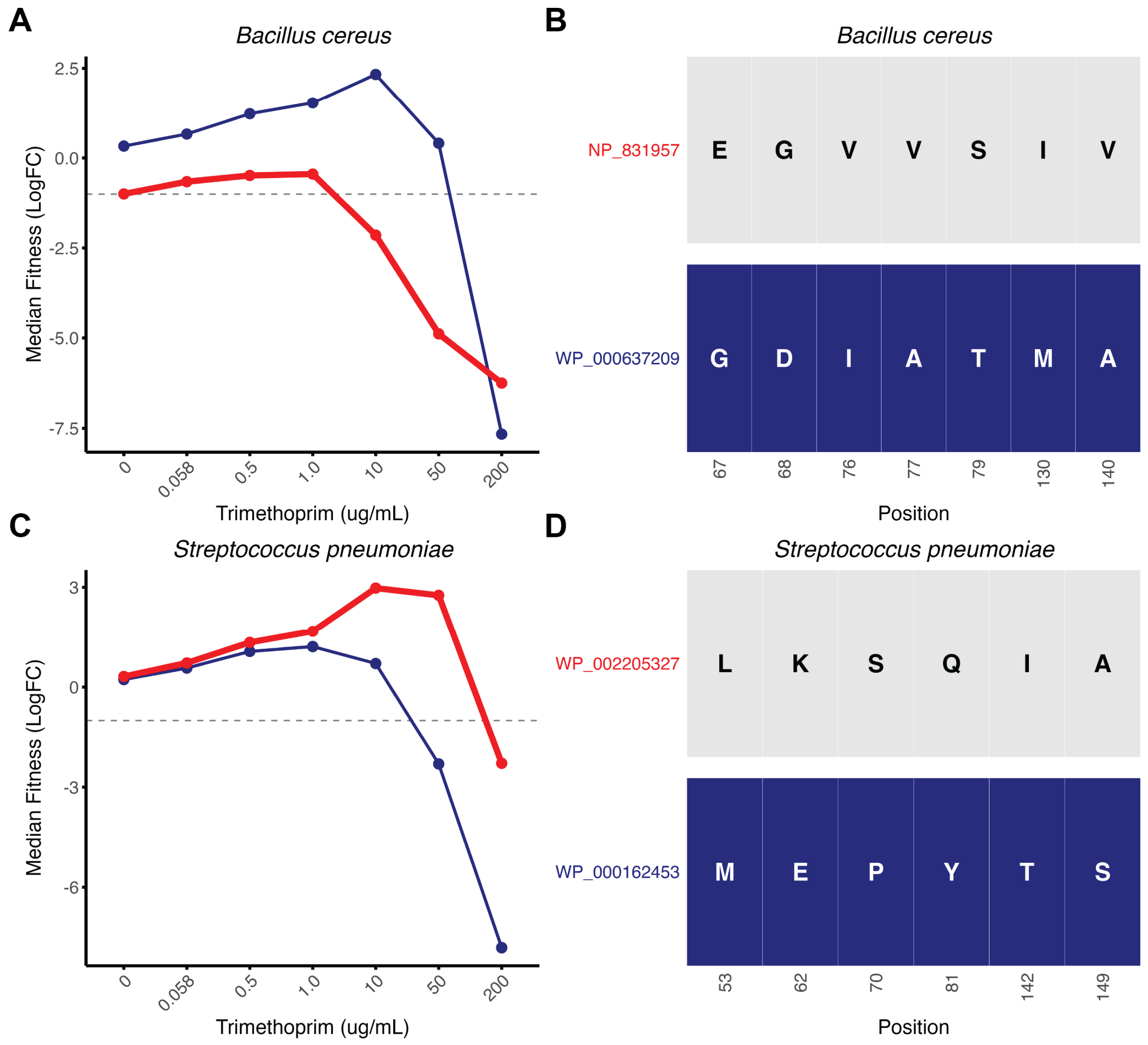

**Fig. S17. Pathogenic DHFR Homologs and Dial-out Variants.** (**A**) Median fitness of *Bacillus cereus* across the TMP gradient, comparing the reference variant (NP_831957; red line) for the resistance analysis with the dial-out variant (WP_000637209; blue line) used for fitness validation. (**B**) Grid plot of amino acid substitutions in *B. cereus* (NP_831957; top row) showing 7 unique residue substitutions in the dial-out variant (WP_000637209; bottom row). (**C**) Median fitness of *Streptococcus pneumoniae* across the TMP gradient, comparing the reference variant (WP_002205327; red line) for the resistance analysis with the dial-out variant (WP_000162453; blue line) used for fitness validation. (**D**) Grid plot of amino acid substitutions in *S. pneumoniae* (WP_002205327; top row) showing 6 unique residue substitutions in the dial-out variant (WP_000162453; bottom row).

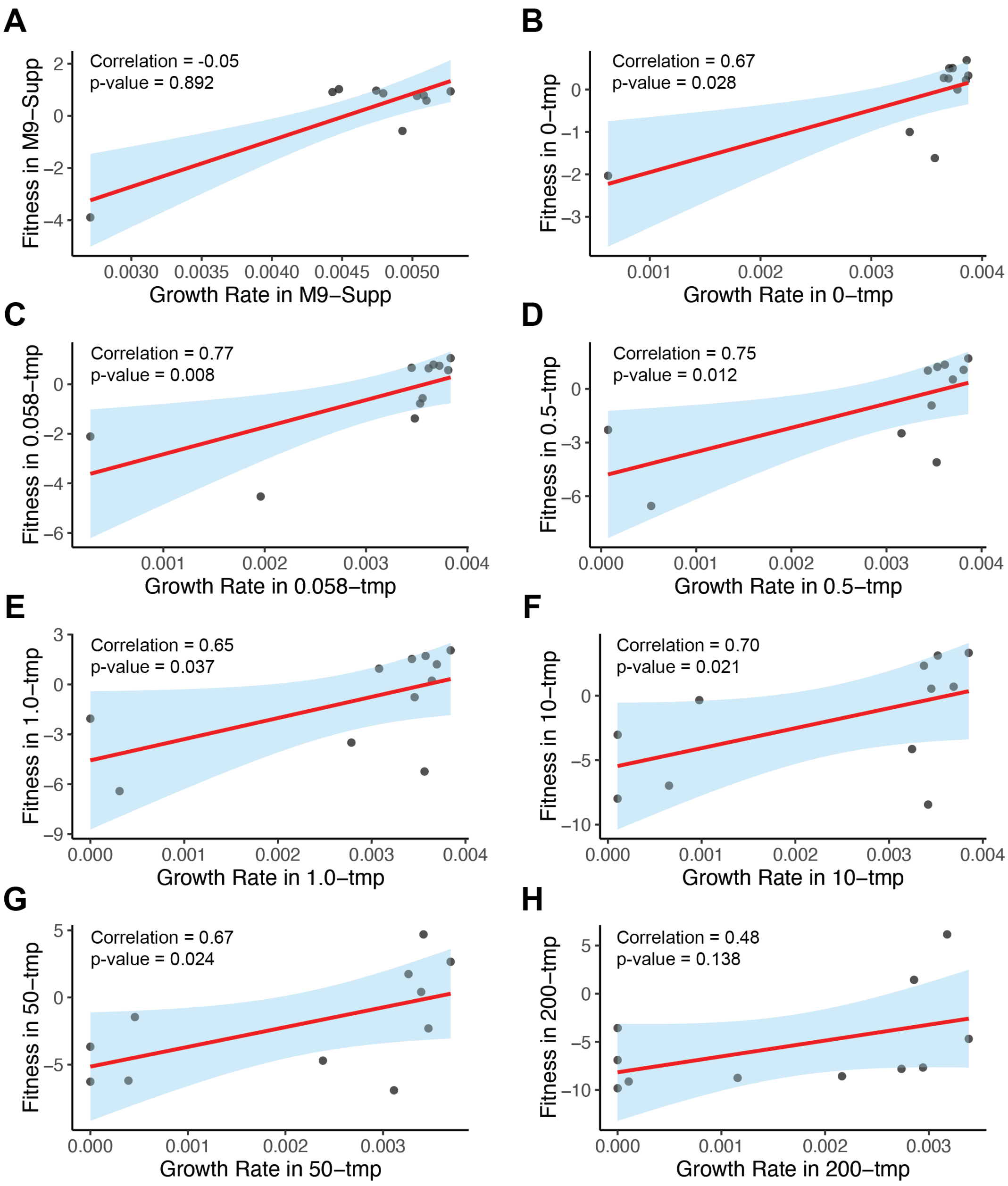

**Fig. S18. Correlation Between Pooled Fitness and Dial-out Growth Rates.** Growth rates of individual homologs, measured via dial-out PCR and plate reader assays, are compared to corresponding fitness values from pooled assays. Spearman's correlation ($r_s$, red line) with a 95% confidence interval (blue ribbon) is shown for: (**A**) M9-Supp media, (**B**) M9 media without TMP (Complementation), (**C**) M9 media with 0.058 µg/mL TMP, (**D**) M9 media with 0.5 µg/mL TMP (MIC), (**E**) M9 media with 1.0 µg/mL TMP, (**F**) M9 media with 10 µg/mL TMP, (**G**) M9 media with 50 µg/mL TMP, and (**H**) M9 media with 200 µg/mL TMP (400× MIC). Growth rate ($min^{-1}$) is defined as the maximum slope of OD600 vs. time on a log-linear plot. Points represent 10 individual homologs and the wild-type *E. coli* DHFR homolog (positive control) spanning fitness values from strongly depleted (median $log_2$ fold-change = -9.1) to highly resistant (median $log_2$ fold-change = +6.2) under 200 µg/mL trimethoprim.

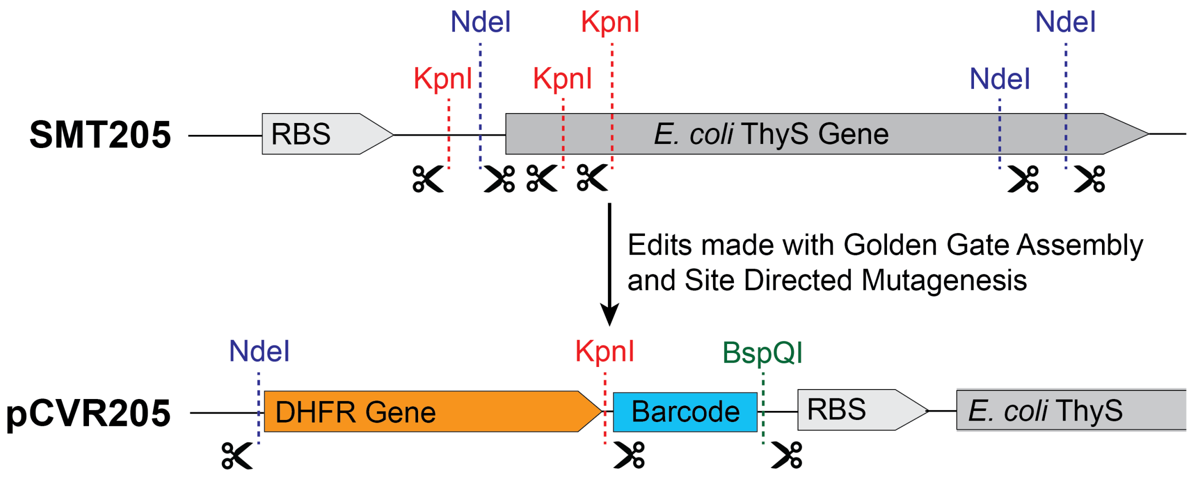

**Fig. S19. pCVR205 Plasmid Construction.** The DHFR expression vector for *E. coli* was derived from the SMT205 plasmid (Addgene #134817). SMT205 was modified to remove unnecessary restriction enzyme sites used for cloning the library. Three restriction sites were then reintroduced to align with the *folA* gene library. To generate the final plasmid, pCVR205, NdeI and KpnI cut sites were removed from the wild-type *thyA* gene, and a gBlock was used to eliminate restriction enzyme sites and insert one KpnI and one BspQI site at the 3' end of the DHFR gene's C-terminus. This was achieved via Golden Gate assembly, integrating the gBlock into the SMT205 backbone.

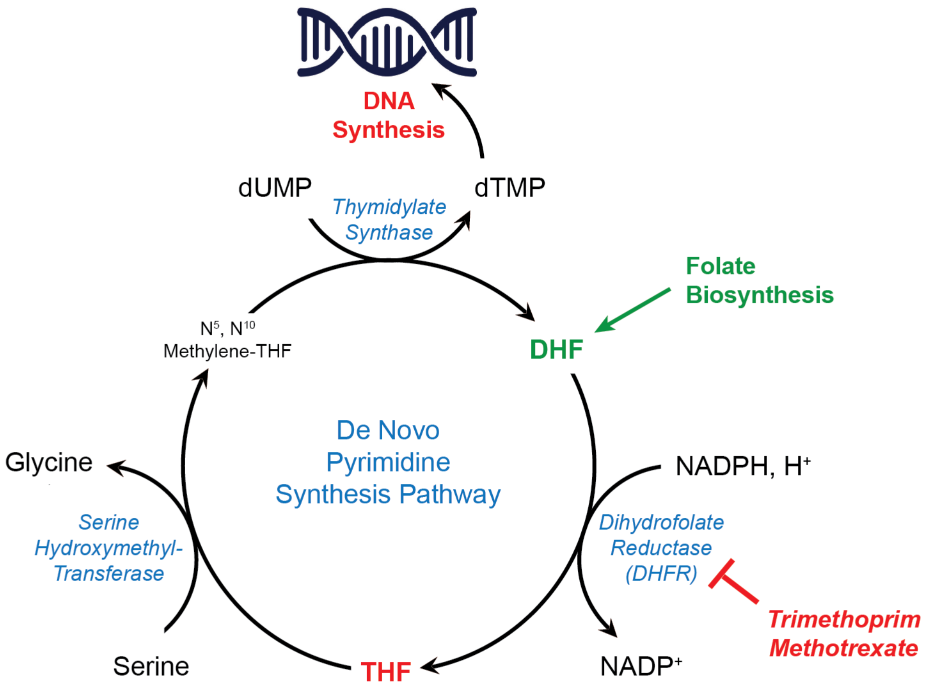

**Fig. S20. Pyrimidine Synthesis.** Dihydrofolate reductase (DHFR) catalyzes the reduction of dihydrofolate (DHF) to tetrahydrofolate (THF) with the cofactor NADPH. DHFR function is tightly coupled with thymidylate synthase. This cycle is an intermediate step in the synthesis of thymidine, one of the four canonical DNA bases. DHFR enzyme activity is inhibited by folate inhibitors such as trimethoprim and methotrexate.

**Table S1. Phylogenetic Summary of DropSynth-Assembled DHFR Homologs.** Phylogenetic summary of 1,208 designed DHFR homologs recovered from the original 1,536 DropSynth library.

| Count | Taxonomy | Count | Domain | Count | % | Phylum |
|---|---|---|---|---|---|---|
| 2 | Domain | 1,188 | Bacteria | 478 | 40.1 | Pseudomonadota |
| 19 | Phylum | 18 | Archaea | 472 | 39.6 | Bacillota |
| 31 | Class | 2 | Virus | 118 | 9.9 | Bacteroidota |
| 74 | Order | | | 61 | 5.1 | Actinomycetota |
| 159 | Family | | | 19 | 1.6 | Mycoplasmatota |
| 330 | Genus | | | 18 | 1.5 | Euryarchaeota (Archaea) |
| 778 | Species | | | 6 | 0.5 | Chlamydiota |
| | | | | 3 | 0.3 | Myxococcota |
| | | | | 3 | 0.3 | Planctomycetota |
| | | | | 3 | 0.3 | Spirochaetota |
| | | | | 3 | 0.3 | Thermotogota |
| | | | | 2 | 0.2 | Uroviricota (Virus) |
| | | | | 1 | 0.1 | Bdellovibrionota |
| | | | | 1 | 0.1 | Candidatus Saccharibacteria |
| | | | | 1 | 0.1 | Cyanobacteria |
| | | | | 1 | 0.1 | Fusobacteriota |
| | | | | 1 | 0.1 | Lentisphaerota |
| | | | | 1 | 0.1 | Thermodesulfobacteriota |
| | | | | 1 | 0.1 | Verrucomicrobiota |

**Table S2. Sequence and Barcode Summary.** Summary of total sequences and barcodes (BC) recovered and mapped across all nine experimental conditions for both codon version libraries.

| Codon Version | Media Condition | Raw Sequence Count | Mapped Sequence Count | Mapped Sequences (%) | Raw Barcode Count | Mapped Barcode Count | Mapped Barcodes (%) |
|---|---|---|---|---|---|---|---|
| Codon 1 | LB | 83,770,481 | 61,036,972 | 73 | 332,234 | 197,558 | 59 |
| Codon 1 | M9: Supplemented | 108,604,061 | 79,808,887 | 73 | 359,595 | 145,597 | 40 |
| Codon 1 | M9: Non-Supplemented (0 ug/mL TMP) | 54,119,812 | 39,528,966 | 73 | 246,911 | 95,345 | 39 |
| Codon 1 | M9: Non-Supplemented (0.058 ug/mL TMP) | 47,847,019 | 34,955,335 | 73 | 174,073 | 87,057 | 50 |
| Codon 1 | M9: Non-Supplemented (0.5 ug/mL TMP) | 40,802,227 | 29,804,232 | 73 | 161,495 | 76,336 | 47 |
| Codon 1 | M9: Non-Supplemented (1.0 ug/mL TMP) | 39,746,862 | 28,977,937 | 73 | 171,737 | 71,182 | 41 |
| Codon 1 | M9: Non-Supplemented (10 ug/mL TMP) | 53,156,432 | 38,534,158 | 72 | 103,600 | 51,972 | 50 |
| Codon 1 | M9: Non-Supplemented (50 ug/mL TMP) | 49,744,711 | 36,333,615 | 73 | 179,548 | 55,599 | 31 |
| Codon 1 | M9: Non-Supplemented (200 ug/mL TMP) | 38,565,223 | 29,013,111 | 75 | 46,363 | 15,161 | 33 |
| **Codon 1** | **Total Counts and Percentage Mapped** | **516,356,828** | **377,993,213** | **73** | **1,775,556** | **795,807** | **45** |
| Codon 2 | LB | 48,821,977 | 34,177,971 | 70 | 428,302 | 266,916 | 62 |
| Codon 2 | M9: Supplemented | 65,871,632 | 45,234,084 | 69 | 409,052 | 191,349 | 47 |
| Codon 2 | M9: Non-Supplemented (0 ug/mL TMP) | 68,405,404 | 47,717,303 | 70 | 291,387 | 141,835 | 49 |
| Codon 2 | M9: Non-Supplemented (0.058 ug/mL TMP) | 43,914,011 | 30,754,811 | 70 | 215,066 | 126,987 | 59 |
| Codon 2 | M9: Non-Supplemented (0.5 ug/mL TMP) | 43,752,442 | 30,367,898 | 69 | 190,020 | 108,523 | 57 |
| Codon 2 | M9: Non-Supplemented (1.0 ug/mL TMP) | 49,596,583 | 34,229,832 | 69 | 181,809 | 101,164 | 56 |
| Codon 2 | M9: Non-Supplemented (10 ug/mL TMP) | 58,144,156 | 38,659,435 | 66 | 151,536 | 76,505 | 50 |
| Codon 2 | M9: Non-Supplemented (50 ug/mL TMP) | 52,530,782 | 33,688,875 | 64 | 134,374 | 67,666 | 50 |
| Codon 2 | M9: Non-Supplemented (200 ug/mL TMP) | 63,525,568 | 39,388,556 | 62 | 170,613 | 66,503 | 39 |
| **Codon 2** | **Total Counts and Percentage Mapped** | **494,562,555** | **334,218,765** | **68** | **2,172,159** | **1,147,448** | **53** |
| **Both** | **Total Counts and Percentage Mapped** | **1,010,919,383** | **712,211,978** | **70** | **3,947,715** | **1,943,255** | **49** |

**Table S3. False-positive Rates.** False-positive rates (fitness ≥ -1) were estimated using highly mutated variants (>50 amino acid substitutions) with a minimum of 5 unique barcodes.

| Experimental Condition | Codon 1 | | | Codon 2 | | |
|---|---|---|---|---|---|---|
| | Total Mutants (mutations > 49) | False-Positive Mutants (fitness > -1) | False-Positive Rate (%) | Total Mutants (mutations > 49) | False-Positive Mutants (fitness > -1) | False-Positive Rate (%) |
| Complementation | 16,239 | 96 | 0.6% | 15,121 | 197 | 1.3% |
| MIC (TMP) | 12,160 | 81 | 0.7% | 11,321 | 159 | 1.4% |
| 400x MIC (TMP) | 1,068 | 13 | 1.2% | 5,222 | 30 | 0.6% |

**Table S4. Homolog Summary by Codon Library.** Summary of unique homologs recovered from the DropSynth-assembled mapping effort, raw sequence counts derived from the complementation assay conditions (min. 1 sequence per barcode), filtered sequence counts derived from the complementation assay conditions (min. 10 sequences per barcode), and filtered sequence counts with a min. 5 unique barcodes per homolog in the complementation assay conditions.

| **Codon Version** | **Mapping Homologs** (DropSynth) | **Raw Count Homologs** (min. 1 seq) | **Filtered Count Homologs** (min. 10 seqs) | **Filtered BC Homologs** (min. 5 BCs) |
|---|---|---|---|---|
| **Codon 1** | 1,048 | 973 | 961 | 797 |
| **Codon 2** | 904 | 840 | 818 | 666 |
| **Shared** | 744 | 663 | 643 | 493 |
| **Unique** | 464 | 487 | 493 | 477 |
| **Total** | 1,208 | 1,150 | 1,136 | 970 |

**Table S5. Resistant Homologs Recovered from 200 ug/mL TMP Condition**. Summary table of the top 16 homologs with sufficient fitness (> -1) to retain function under the highest TMP assay condition. Codon indicates sufficient fitness (> -1) in Codon 1, Codon 2, or both.

| Codon | mutID | fitD11D03 | fitE06D04 | Phylum | Species |
|---|---|---|---|---|---|
| Codon 1 | WP_003686654 | 1.74 | -3.64 | Betaproteobacteria | *Neisseria flavescens* |
| Codon 1 | WP_002839715 | 1.6 | -5.3 | Bacillota | *Finegoldia magna* |
| Codon 1 | WP_003498667 | 0.92 | -3.23 | Bacillota | *NA* |
| Codon 1 | WP_008152545 | -0.58 | -5.39 | Bacteroidota | *Parabacteroides johnsonii* |
| Codon 1 | WP_004442738 | -0.6 | -6.28 | Alphaproteobacteria | *NA* |
| Codon 1 | WP_008237079 | -0.96 | -3.81 | Bacteroidota | *Imtechella halotolerans* |
| Codon 2 | WP_008754633 | -1.91 | -0.59 | Bacillota | *Lachnoanaerobaculum saburreum* |
| Codon 2 | WP_008664019 | -8.5 | 3.75 | Bacteroidota | *NA* |
| Codon 2 | WP_009161670 | -9.93 | 2.08 | Bacteroidota | *Hoylesella saccharolytica* |
| Both | WP_010075211 | 6.62 | 5.48 | Bacillota | *Clostridium cellulovorans* |
| Both | WP_007654866 | 6.48 | 5.41 | Bacteroidota | *NA* |
| Both | WP_008990832 | 4.94 | 1.96 | Bacteroidota | *Galbibacter marinus* |
| Both | WP_008583481 | 2.65 | 1.78 | Bacteroidota | *Niabella soli* |
| Both | WP_009778768 | 2.3 | 1.48 | Bacteroidota | *Leeuwenhoekiella blandensis* |
| Both | NP_229441 | -0.55 | -0.77 | Thermotogota | *Thermotoga maritima* |
| Both | WP_008390643 | -0.71 | 0.21 | Bacillota | *NA* |

**Table S6. Resistant Homolog Summary Across TMP Gradient.** Summary of the number of perfect assembly homologs resistant to trimethoprim (fitness > -1) across the TMP gradient. The total homolog count varies between TMP treatments based on recovery rates.

| Homolog Fitness | 0 ug/mL TMP | 0.058 ug/mL TMP | 0.5 ug/mL TMP | 1.0 ug/mL TMP | 10 ug/mL TMP | 50 ug/mL TMP | 200 ug/mL TMP |
|---|---|---|---|---|---|---|---|
| Complement (fit > -1) | 416 | 318 | 246 | 226 | 128 | 80 | 27 |
| Dropout (fit < -1) | 0 | 92 | 162 | 183 | 276 | 321 | 332 |
| Total Homologs | 416 | 410 | 408 | 409 | 404 | 401 | 359 |

**Table S7. Phylogenetic Fitness Differences Across TMP Gradient.** Summary statistics for absolute differences in fitness between homolog pairs based on evolutionary distance for each TMP treatment. Close-related pairs have an absolute difference in evolutionary distance < 1, whereas far-related pairs have an absolute difference in evolutionary distance > 1. Fitness differences are calculated as the absolute value of the difference between fitness values of two taxa, resulting in a unitless quantity.

| Treatment | Group | N | Mean | SD | Min | Max |
|---|---|---|---|---|---|---|
| 0.058 | Close | 5,814 | 0.70 | 0.79 | 0 | 5.87 |
| | Far | 161,876 | 1.50 | 1.51 | 0 | 9.05 |
| 0.5 (MIC) | Close | 5,812 | 1.48 | 1.40 | 3.44E-04 | 8.65 |
| | Far | 160,244 | 2.71 | 2.32 | 6.28E-06 | 10.08 |
| 1 | Close | 5,812 | 1.88 | 1.70 | 3.06E-04 | 9.29 |
| | Far | 161,060 | 3.09 | 2.48 | 1.67E-05 | 10.22 |
| 10 | Close | 5,744 | 3.68 | 3.19 | 2.23E-03 | 12.79 |
| | Far | 157,068 | 4.31 | 3.04 | 0 | 13.69 |
| 50 | Close | 5,764 | 2.88 | 2.57 | 3.21E-04 | 11.42 |
| | Far | 154,636 | 3.34 | 2.62 | 2.12E-05 | 13.40 |
| 200 (400x MIC) | Close | 5,254 | 2.24 | 2.40 | 2.60E-04 | 15.54 |
| | Far | 123,268 | 3.14 | 3.43 | 0 | 18.20 |

**Table S8. Dial-out Variant Pooled Fitness Summary.** Median fitness for pooled DHFR dial-out variants across the trimethoprim (TMP) gradient.

| mutID | M9-Supp | 0-TMP | 0.058-TMP | 0.5-TMP | 1.0-TMP | 10-TMP | 50-TMP | 200-TMP |
|---|---|---|---|---|---|---|---|---|
| NP_267306 | -0.58 | -1.62 | -1.38 | -0.93 | -0.77 | 0.55 | 1.74 | 1.44 |
| WP_000637209 | 0.58 | 0.33 | 0.66 | 1.23 | 1.53 | 2.33 | 0.41 | -7.66 |
| WP_008976421 | 0.94 | 0.50 | 0.79 | 1.02 | 0.95 | -0.34 | -1.46 | -6.89 |
| WP_003776922 | -3.89 | -2.03 | -2.11 | -2.29 | -2.06 | -3.03 | -3.67 | -3.56 |
| WP_008578924 | 0.97 | 0.27 | 0.64 | 1.35 | 1.72 | 3.12 | 4.70 | 6.16 |
| WP_002897636 | 1.02 | 0.69 | 1.05 | 1.70 | 2.05 | 3.33 | 2.66 | -4.69 |
| WP_000162453 | 0.86 | 0.22 | 0.57 | 1.06 | 1.21 | 0.70 | -2.31 | -7.81 |
| WP_003012456 | 0.79 | 0.00 | -0.56 | -2.48 | -3.50 | -6.98 | -6.20 | -9.12 |
| WP_000162462 | 0.91 | 0.51 | 0.75 | 0.53 | 0.23 | -4.14 | -4.71 | -8.74 |
| WP_003027976 | 0.76 | 0.26 | -0.78 | -4.11 | -5.24 | -8.45 | -6.92 | -8.57 |
| WT-DHFR | NA | -1.00 | -4.53 | -6.54 | -6.42 | -7.99 | -6.27 | -9.82 |
| mCherry | NA | -1.29 | -1.57 | -1.75 | -1.54 | -2.26 | -2.16 | -4.44 |

**Table S9. Dial-out Variant Taxonomic Summary.** Phylogenetic taxonomic classification for each dial-out variant and percent sequence identity to *E. coli*. IDs in **bold** correspond to the pathogenic homologs *Bacillus cereus* and *Streptococcus pneumoniae*.

| mutID | Sequence Identity to E. coli (%) | Phylum | Class | Order | Family | Genus | Species |
|---|---|---|---|---|---|---|---|
| NP_267306 | 45.2 | Bacillota | Bacilli | Lactobacillales | Streptococcaceae | Lactococcus | Lactococcus lactis |
| **WP_000637209** | **45.7** | **Bacillota** | **Bacilli** | **Bacillales** | **Bacillaceae** | **Bacillus** | **Bacillus cereus** |
| WP_008976421 | 45.1 | Bacillota | Clostridia | Lachnospirales | Lachnospiraceae | *NA* | Lachnospiraceae bacterium |
| WP_003776922 | 44.5 | Bacillota | Bacilli | Lactobacillales | Carnobacteriaceae | Alloiococcus | Alloiococcus otitis |
| WP_008578924 | 38.1 | C. Saccharibacteria | *NA* | *NA* | *NA* | *NA* | candidate division TM7 genomosp. GTL1 |
| WP_002897636 | 44.1 | Bacillota | Bacilli | Lactobacillales | Streptococcaceae | Streptococcus | Streptococcus sanguinis |
| **WP_000162453** | **44.0** | **Bacillota** | **Bacilli** | **Lactobacillales** | **Streptococcaceae** | **Streptococcus** | **Streptococcus pneumoniae** |
| WP_003012456 | 50.6 | Bacteroidota | Sphingobacteriia | Sphingobacteriales | Sphingobacteriaceae | Sphingobacterium | Sphingobacterium spiritivorum |
| WP_000162462 | 44.0 | Bacillota | Bacilli | Lactobacillales | Streptococcaceae | Streptococcus | Streptococcus mitis |
| WP_003027976 | 41.8 | Bacillota | Bacilli | Lactobacillales | Streptococcaceae | Streptococcus | *NA* |

**Table S10. Primers for Plasmid Preparation.** Primer pairs for plasmid preparation and gBlock sequence designed to insert one KpnI site and one BspQI site at the 3′ end of the DHFR gene. This gBlock was integrated into the SMT205 backbone using Golden Gate assembly, creating the plasmid pCVR1.

| Primer Pair Name | Forward Sequence (5’-->3’) | Reverse Sequence (5’-->3’) |
|---|---|---|
| pCVR205_SDM | tatgatcagtctgattgcggcgttagcggtagatcgcgttaTCGGCATGGAAAACGCCA | tgatatctcattattaaagttaaacaaaattatttctacaggGGAATTGTTATCCGCTCACAATTC |
| SMT205_bb | TGACCAGGTCTCGGCAATTGAGCCGTGAGCCG | TGACCAGGTCTCGCCAGAATCTCAAAGCAATAGCTGTGA |
| mi3_R1 | GTGGAATTGTGAGCGGATAACAATTTCACACAGGAAACAGCTCATATG | *NA* |
| EV4_BspQI | *NA* | cagtctaGCTCTTCatgaCGCACATTTCCCCGAAAAGTGCCACCTGACGTCg |

| gBlock Sequence from IDT |
|---|
| TGACCAGGTCTCGCTGGAGCGGCGGTAAGGTACCTAAGTGTGGCTGCGGAACGCA<br>CGACGTCAGGTGGCACTTTTCGGGGAAATGTGCGTCATGAAGAGCCAGGCGTCGAC<br>AAGCTTGCGGCCGCATAATGCTTAAGTCGAACAGAAAGTAATCGTATTGTACATCCC<br>TATCAGTGATAGAGATTGACATCCCTATCAGTGATAGAGATACTGAGCACATCAGCA<br>GGACGCACTGACCGAATTCATTAAAGAGGAGAAAGGGACCACATGGCAGATCTCAT<br>GAAACAGTACCTGGAGCTGATGCAAAAAGTTCTGGATGAGGGGACCCAGAAAAACG<br>ACCGCACGGGGACCGGAACGCTGAGCATTTTTGGCCATCAGATGCGCTTTAACCTG<br>CAGGACGGATTCCCGCTGGTTACGACCAAACGCTGCCACCTGCGTAGCATTATTCA<br>TGAGCTGCTGTGGTTTCTGCAAGGTGACACTAACATCGCGTATCTGCACGAAAACAA<br>TGTGACGATCTGGGATGAATGGGCCGATGAAAACGGCGATCTGGGCCCAGTGTATG<br>GTAAACAGTGGCGCGCCTGGCCAACGCCAGATGGCCGCCACATCGATCAGATCAC<br>GACCGTGCTGAATCAACTGAAAAACGACCCGGACAGCCGCCGCATTATTGTTTCCG<br>CGTGGAATGTGGGTGAACTGGATAAAATGGCGCTGGCGCCGTGCCATGCATTCTTC<br>CAGTTTTACGTTGCGGACGGTAAGCTGAGCTGTCAACTTTATCAGCGCAGCTGCGAT<br>GTTTTCCTCGGCCTGCCGTTCAATATCGCCAGCTACGCGTTACTGGTGCACATGATG<br>GCGCAGCAGTGCGACCTGGAGGTGGGTGATTTTGTCTGGACCGGCGGTGATACCCA<br>CCTGTACAGCAACCACATGGACCAAACGCATCTGCAACGAGACCTCGTCA |

**Table S11. iTag Illumina Prep Primers.** PCR was performed to amplify plasmid barcodes using five primer pairs that introduced custom sequencing adapters and library indexes compatible with Illumina sequencing.

| Primer Pair Name | Forward Sequence (5'-->3') | Reverse Sequence (5'-->3') |
|---|---|---|
| CVR205stubBC1_FWD | GCTCTTCCGATCTNGGTACCtaaGTGTGGCTGCGGAAC | GCTCTTCCGATCTNGTGCCACCTGACGTCgtgc |
| CVR205stubBC2_FWD | GCTCTTCCGATCTNNGGTACCtaaGTGTGGCTGCGGAAC | GCTCTTCCGATCTNNGTGCCACCTGACGTCgtgc |
| CVR205stubBC3_FWD | GCTCTTCCGATCTNNNGGTACCtaaGTGTGGCTGCGGAAC | GCTCTTCCGATCTNNNGTGCCACCTGACGTCgtgc |
| CVR205stubBC4_FWD | GCTCTTCCGATCTNNNNGGTACCtaaGTGTGGCTGCGGAAC | GCTCTTCCGATCTNNNNGTGCCACCTGACGTCgtgc |
| CVR205stubBC5_FWD | GCTCTTCCGATCTNNNNNGGTACCtaaGTGTGGCTGCGGAAC | GCTCTTCCGATCTNNNNNGTGCCACCTGACGTCgtgc |

**Table S12. Dial-out PCR Primers.** Dial-out PCR primers were designed to flank each homolog construct, with reverse primers annealing to the gene-specific barcode.

| mutID | Primer Pair | Barcode | Forward Sequence (5'-->3') | Reverse Sequence (5'-->3') |
|---|---|---|---|---|
| NP_267306 | NP_06_FWD NP_06_REV | ACCCGGCGTGGATATATCTA | GATATCATATGATCATTGGCATCTGGG | TAGATATATCCACGCCGGGT |
| WP_000637209 | WP_09_FWD WP_09_REV | AAACCGAATTTTTGCATGGA | GATATCATATGATCGTGAGCTTTATGGTT | GCTCCATGCAAAAATTCGGTTT |
| WP_008976421 | WP_21_FWD WP_21_REV | AAAGCGTCGTGTAAGCGATC | GATATCATATGAACATTATTGTCGCGGTTG | GATCGCTTACACGACGCTTT |
| WP_003776922 | WP_22_FWD WP_22_REV | ACTCGACCTCTAAAAATCTT | GATATCATATGATTGCCTACGTTTGG | GCAAGATTTTTAGAGGTCGAGT |
| WP_008578924 | WP_24_FWD WP_24_REV | TACACTGCCCTGAATTATCT | GATATCATATGAAAGTGTTTCTGATTGTCGC | GCAGATAATTCAGGGCAGTGTA |
| WP_002897636 | WP_36_FWD WP_36_REV | CCAGTAGGAGCAAAGTCTAC | GATATCATATGACGAAAAAAATCATTGCA | GTAGACTTTGCTCCTACTGG |
| WP_000162453 | WP_53_FWD WP_53_REV | CATCTTAGTCCTCGATTTAT | GATATCATATGACCAAAAAGATTGTCGC | GTGCATAAATCGAGGACTAAGATG |
| WP_003012456 | WP_56_FWD WP_56_REV | ATCGCACGGGTCTGCGCCGA | GATATCATATGTCGCACCTGAAAATCAC | CATACACGCACTACCCCTGT |
| WP_000162462 | WP_62_FWD WP_62_REV | AGTGGGCATTAGTATAGATT | GATATCATATGACTAAAAAAATTGTGGCG | GCAATCTATACTAATGCCCACT |
| WP_003027976 | WP_76_FWD WP_76_REV | CAACTTAGTTTTGAATTGGA | GATATCATATGATTAAAAAAATTGTGGCGAT | GTGCTCCAATTCAAAACTAAGTTG |
